# The evolution of partner specificity in mutualisms

**DOI:** 10.1101/2021.12.10.472103

**Authors:** Christopher Carlson, Erol Akçay, Bryce Morsky

**Affiliations:** Department of Biology, University of Pennsylvania, Philadelphia, PA, USA

**Keywords:** Mutualisms, Symbiosis, Cooperation, Partner Specificity, Generalism, Dispersal

## Abstract

Mutualistic species vary in their level of partner specificity, which has important evolutionary, ecological, and management implications. Yet, the evolutionary mechanisms which underpin partner specificity are not fully understood. Most work on specialization focuses on the trade-off between generalism and specialism, where specialists receive more benefits from preferred partners at the expense of benefits from non-preferred partners, while generalists receive similar benefits from all partners. Because all mutualisms involve some degree of both cooperation and conflict between partners, we highlight that specialization to a mutualistic partner can be cooperative, increasing benefit to a focal species and a partner, or antagonistic, increasing resource extraction by a focal species from a partner. We devise an evolutionary game theoretic model to assess the evolutionary dynamics of cooperative specialization, antagonistic specialization, and generalism. Our model shows that cooperative specialization leads to bistability: stable equilibria with a specialist host and its preferred partner excluding all others. We also show that under cooperative specialization with spatial effects, generalists can thrive at the boundaries between differing specialist patches. Under antagonistic specialization, generalism is evolutionarily stable. We provide predictions for how a cooperation-antagonism continuum may determine the patterns of partner specificity that develop within mutualistic relationships.

## 1 Introduction

Mutualisms enhance the biodiversity in global ecosystems (Bascompte, 2019); therefore, understanding the evolutionary underpinnings of such associations is crucial to conservation and or promotion of biodiversity at the local, regional, and global scale (Bronstein et al., 2004). One crucial factor in the development of complex mutualism is partner specificity, the degree to which one organism preferentially interacts with a single partner-species (Chomicki et al., 2020), which has been examined in numerous systems in which partners are acquired horizontally from the surrounding environment including legume-Rhizobium, fig-wasp, and damselfish-sea anemone mutualisms (Wang et al., 2012; Bronstein, 1987). The prevalence of partner specificity within horizontally transmitted interactions is difficult to explain given that there is a fitness cost associated with a mismatch between a host and its symbiont (Uchiumi and Sasaki, 2020; Batstone et al., 2020). Because specificity plays a pivotal role in shaping the stability, niche, and coevolutionary dynamics of mutualisms (Chomicki et al., 2020; Harrison et al., 2018; Uchiumi and Sasaki, 2020), variation in partner specificity may be explained by incorporating partner cooperation into a theoretical framework to examine how specificity and cooperation co-evolve in horizontally transmitted partnerships.

There is significant variability in the degree to which partners cooperate across both general and specific horizontally transmitted mutualisms such as plant-pollinator, coral-algae, and seed dispersal mutualisms (Gomulkiewicz et al., 2003; Hoeksema and Bruna, 2000; Bogdziewicz et al., 2018; Stat et al., 2008) which likely shapes the evolutionary stability of generalism vs. specialism. Some symbiotic species display little partner preference, forming partnerships with a wide variety of partner species (e.g. ant-plant and plant-mycorrhizal mutualisms involve generalist partners (Chomicki and Renner, 2017; Peay et al., 2015)) indicating that in the absence of one partner, another may serve as a substitute. Others are highly specific and only interact with as few as one other mutualistic partner species (e.g. moth-yucca plant, wasp-fig) (Gomulkiewicz et al., 2003; Machado et al., 2005). In such interactions, absence of a preferred partner could result in death. It is likely that specific mutualisms involve a high degree of pairwise co-evolution, maximizing benefits when preferred partners interact, while reducing the benefits derived from associating with other partners. This trade-off makes specialism an apparently risky strategy relative to generalism, as absence/low abundance of a preferred partner may be highly detrimental to the fitness of the specialist (Thrall et al., 2007).

There is another and relatively less explored dimension to the generalist-specialist conundrum. Many mutualisms involve exchange of benefits that are costly to produce, and therefore even when the net outcome is mutually beneficial, there is an underlying conflict of interest. This conflict of interest may be resolved through behavioral mechanisms such as negotiation (Akçay and Roughgarden, 2007), or evolutionarily through adaptations and counter adaptations (as in nectar robbing, Irwin et al., 2010). The resolution mechanisms determine the division of benefits from an interaction, and may cause one partner benefit more at the expense of the other. At the same time, mutualisms often also involve intricate behavioral adaptations and physiological adaptations to produce the benefits that both partners benefit from. Mutual specialization in these adaptations can lead to production of more benefits for both partners. This mutual specialization is reinforced by partner fidelity feedback given that the partners that cooperate the most receive the greatest benefit from mutualism (Bull and Rice, 1991; Sachs et al., 2004; Friesen, 2012). Physiological and behavioral adaptations mediate this cooperation in ant-plant mutualisms in which host plant capacity to house ants in domatia is linked to ant colony protection from herbivory (Archetti et al., 2011; Mayer et al., 2014). Thus, from the perspective of one of the partner species we can envision two kinds of specialism: cooperative specialism where both the focal species and its preferred partner enjoy higher benefits relative to non-preferred partners, and antagonistic specialists, where the focal species gets more benefits from its preferred partner, but the preferred partner gets less. These two kinds of specialisms will generate different kinds of ecological feedback (Bever et al., 1997), which will affect the evolutionary conditions under which specialism or generalism prevails.

An example of a system where such dynamics are expected is coral-algal symbioses, which are crucial to marine biodiversity. Within this symbiosis there exists significant variation in partner specificity; some coral hosts specialize, associating closely with a small range of Symbiodinaceae, while other hosts generalize, associating with many diverse symbionts (Thompson and Pellmyr, 1992). There is significant variability in how specific coral-algal associations are (Baker, 2003), yet theoretical models of coral symbiosis have not yet attempted to explain this variation (Raharinirina et al., 2017; Cunning et al., 2017; Roughgarden, 1975). Furthermore, while coral symbioses have been traditionally viewed as cooperative (Muscatine and Porter, 1977), there are also elements of Cnidarian symbiosis that showcase the regulation of conflict in the interaction, such as the coral host limiting the resource availability, reproductive capability, and resource extraction capability of their symbionts (Xiang et al., 2020; Wooldridge, 2010; Sutton and Hoegh-Guldberg, 1990).

Another example system is mycorrhizal symbioses, where generalism is common (Toju et al., 2013; Peay et al., 2015). It is possible that this trend toward generalism is dictated by the fact that dispersal is more difficult in terrestrial systems relative to aquatic systems (Kinlan and Gaines, 2003). Thus, the likelihood of failing to interact with a preferred symbiont is greater. Yet, a few taxa form highly specific associations (Bruns et al., 2002; Sepp et al., 2019) in which hosts and symbionts associate together in strict, pairwise partnerships. Bever (2002) experimentally demonstrated the potential for antagonistic specialization in our sense using plant species *Plantago lanceolata* and *Panicum sphaerocarpon,* where the arbuscular mycorrhizal species growing best with one plant result in poor growth of that plant, and vice versa. At the same time, plant-mycorrhizal symbiosis also presents scope for cooperative specialization in which greater host capability to create a hospitable root environment for particular fungi and greater fungal ability to inhabit this environment likely increases symbiotic benefit for both partners (Hoeksema, 2010).

Antagonistic or cooperative specialization may also underlie mutualisms that resemble domestication. Such interactions often resemble a monopoly in which a host controls a resource, and specializes in a partner that provides an exchange rate favorable to the host, with examples including fungus-growing ants (Formicidae: Attini) and decomposing fungi (family Lepiotaceae) (Villesen et al., 2004), or between damselfish *Tegastes nigricans* and filamentous algae (Hata et al., 2010). In fungus farming ants, for example, increased ant capability to harvest fungi does not necessarily benefit fungal fitness in certain lineages (Shik et al., 2016), representing a potential case of antagonistic specialization. Yet it is also plausible that in many of these interactions mutual behavioral and physiological integration of the partners represent coordinated ways of producing higher mutual benefits.

Here we consider the evolution of specificity using a model in which one partner species can exhibit cooperative or antagonistic specificity on different strains of the other partner species. Specialists in both cases receive an added benefit from a preferred symbiont type at the expense of less benefit with the opposite type. We also consider generalists who receive equal and intermediate benefit from either partner strain. When interacting with their preferred partners, specialists either enhance their partner’s fitness (cooperative specialization), or reduce it (antagonistic specialization) relative to the partner’s interaction with a generalist. We consider the ecological dynamics of these systems in well-mixed and spatially structured populations as well as the evolution of both kinds of specialism as continuous traits, and derive conditions for the evolutionary stability of each given the ecological dynamics they produce. Finally, we analyze how incorporation of population spatial structure may impact the stability of specificity vs. generalism in an evolving population.

## 2 Model

We consider a population of two partner species, which we call “Hosts” and “Symbionts,” for ease of referral to each side, although our model construction applies equally well to non-symbiotic mutualisms such as plant-pollination mutualisms. We assume a simple interaction structure that can represent many different kinds of mutualisms based on the exchange of resources or services (see Figure 1 for a view of the interaction from the host perspective, and Figure 2 for the symbiont perspective). Specifically, we assume that hosts translocate some proportion *x* of a resource, such as inorganic carbon, to their symbiont partner. This partner produces a good or a service, such as organic carbon, which is *γ* times as useful as the original good. The degree of phenotypic matching between hosts and symbionts improves this benefit of symbiosis for both partners as the match between flower and proboscis morphology does in plant-pollinator mutualisms. Symbionts then translocate a proportion *α* of the good they have produced back to their host at the expense of their own resource pool. In this model, hosts and symbionts are in conflict over *α*, signifying the division of benefits, but have mutual interest over increasing *γ*, which increases total benefit from the interaction.

**Figure 1.**
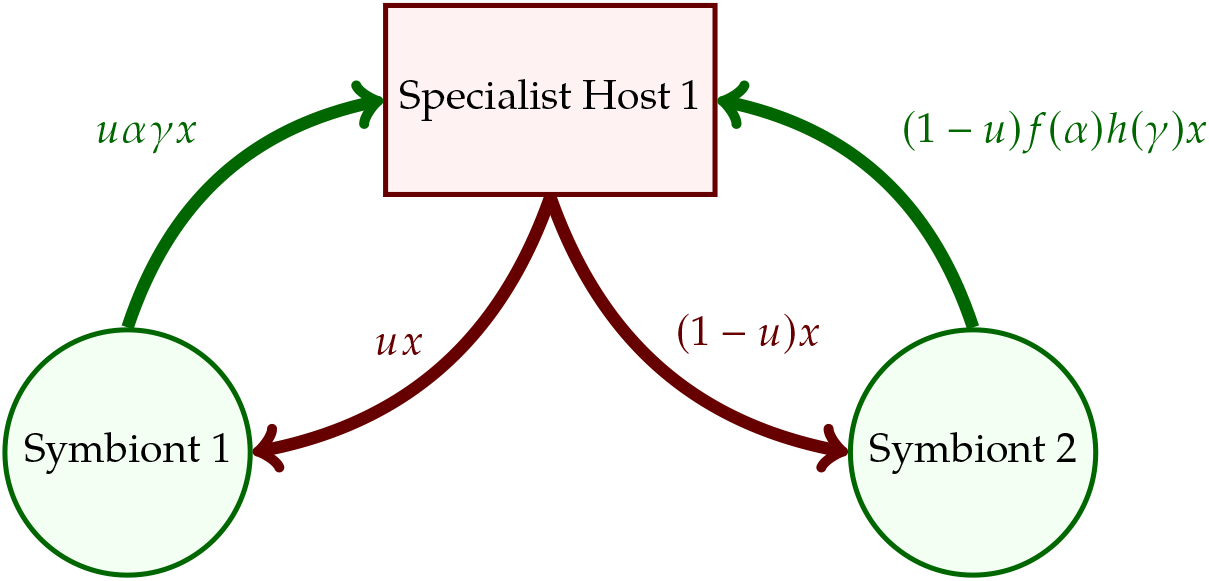
**Associations for specialist host 1. Red arrows denote the amount of the resource given to each symbiont by a focal specialist host 1 with** 0 ≤ *u* ≤ 1. **The symbionts each produce a good or service with value** *γx* **or** *h*(*γ*)*x*. **Green arrows denote the translocation of proportion** *α* **or** *f*(*α*) **of this resource back to the host. Parameters** *α* **and** *γ* **are associated with phenotypes specialized to one another, i.e. those that are “matched”**. *f*(*α*) **and** *h*(*γ*) **are for mismatched mutualisms**.

**Figure 2.**
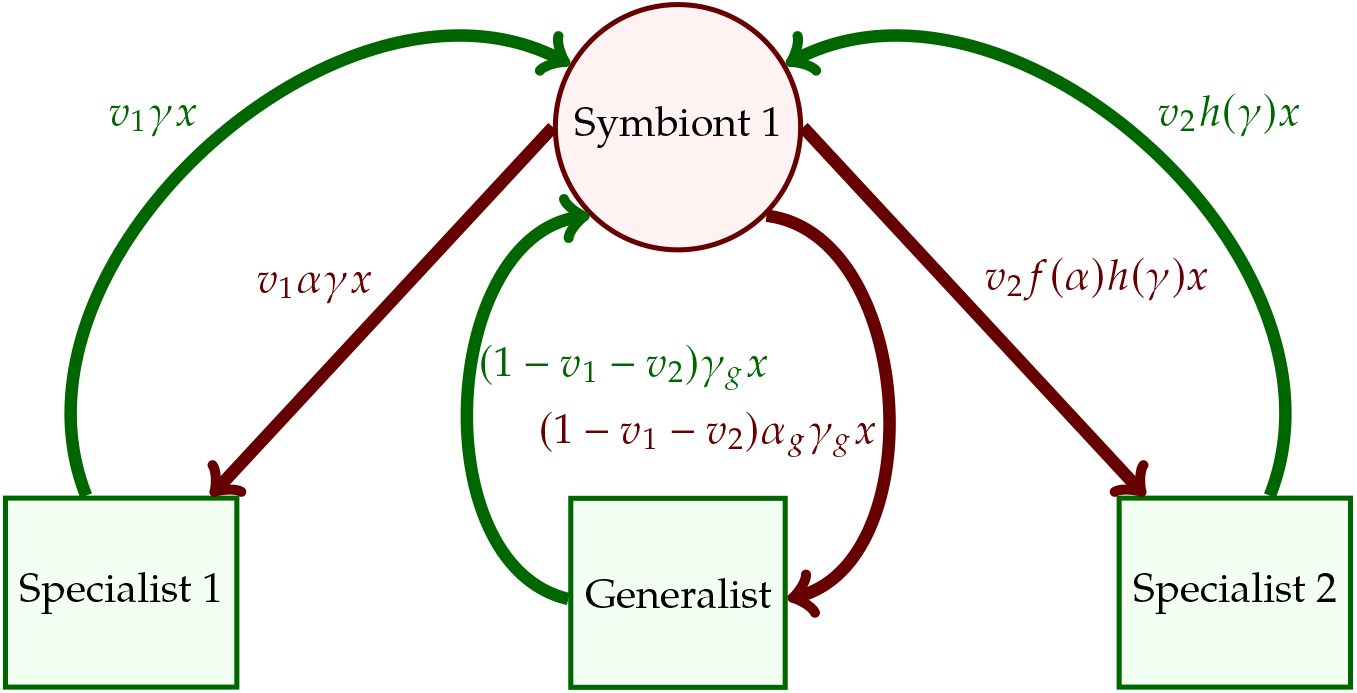
**Associations for symbiont 1. Green arrows denote the value of the resource shared to symbionts by hosts with** 0 ≤ (*v*_1_ + *v*_2_) ≤ 1**. Symbionts produce a good with value** *γx, γ_g_x*, or *h*(*γ*)*x*. **The red arrows indicate the value of the good shared by symbionts to hosts with parameters** *γ_g_* **and** *α_g_* **associated with the generalist host phenotype.**

To allow for the possibility of specialization, we assume that there are two types that make up the symbiont population. Conversely, the host population is made up of three types: hosts that specialize on type 1 or 2 symbionts, and a generalist host. The potential associations for a focal type 1 specialist host are shown in Figure 1, where *u* and 1 – *u* are the frequencies of symbionts 1 and 2, respectively. All symbionts can associate with all host types, but experience and provide different costs and benefits from these associations. Specifically, we assume that all hosts translocate the resource indiscriminately at a cost *x*^2^ to themselves (meaning that the resource is more valuable to the host the less it keeps). How much the hosts get in return from each symbiont depend on the parameters *α* and *γ*. Associations between matching host and symbiont types (e.g. symbiont 1 associates with specialist host 1) correspond to parameters *α* and *γ*, while host and symbiont mismatches (e.g. symbiont 2 associates with specialist host 1) correspond to *f*(*α*) and *h*(*γ*). *f*(*α*) and *h*(*γ*) are trade-off functions in which increasing the value with a preferred type reduces the mismatched value (i.e. *f*′(*α*) < 0 and *h*′(*γ*) < 0). We consider concave up (positive second derivative), concave down (negative second derivative) and linear (second derivative is equal to zero) trade-off functions. Finally, both symbionts associating with generalist hosts exhibit parameters *α_g_* and *γ_g_*. The host payoffs from exclusively interacting with a single symbiont type are:

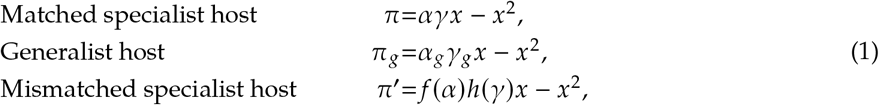

where *π* and *π*′ are the payoffs to specialist hosts matched and mismatched to their preferred symbiont, respectively. (In our symmetric setup, payoffs for specialist host depend only one whether they are matched or mismatched, and not on the identity of the strain they specialize in.) *π_g_* is the generalist host’s payoff, which they earn regardless of which symbiont they are with. We define our payoffs such that matched specialist payoff is greater than generalist specialist payoff, which is greater than the mismatched specialist payoff *π*′, i.e. *π* > *π_g_* > *π*′.

Symbionts associate with one of the three host types where *υ*_1_ is the frequency of host 1, *υ*_2_ is the frequency of host 2, and 1 – *υ*_1_ – *υ*_2_ is the frequency of the generalist host. The payoffs associated with these interactions for one symbiont type are shown in Figure 2. The symbiont payoffs are:

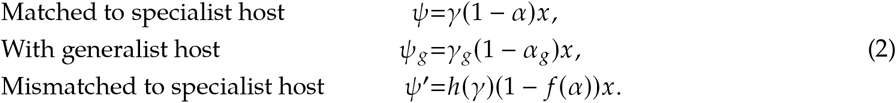

Here, *ψ* and *ψ*′ are the payoffs to symbionts matched and mistmatched to the host that prefers them, respectively, and *ψ_g_* is the payoff to either symbiont when matched with the generalist host. By our assumptions, *ψ* > *ψ_g_* > *ψ*′.

We average these payoffs over the distribution of partners in the population to determine the expected payoffs, 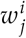. Superscript *i* = *h,s* denotes a host or symbiont, respectively, and subscript *j* = 1,2 denotes type 1 or type 2, respectively. For example, the average payoff for a type 1 host is

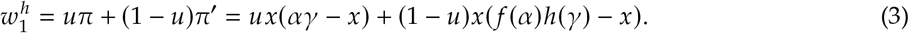

To study the dynamics of the frequencies of hosts and symbionts, we use the replicator equation (Taylor and Jonker, 1978) with two populations, where the change in frequency of a host or symbiont type is determined by the difference between its fitness and the mean fitness of its competitors. Our dynamical equations are thus:

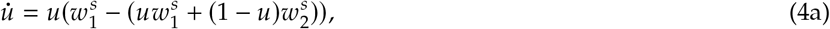

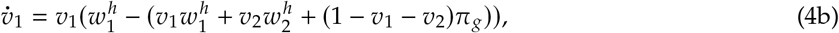

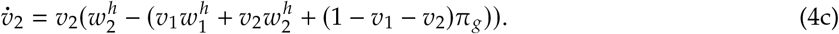

### 2.1 Cooperative and antagonistic specialization

We consider two variations of this general model where the payoffs from specialism are determined in either a cooperative or antagonistic fashion. In the cooperative case, specialist hosts have greater efficiency in their association when they match with their preferred symbiont. We model this as the fraction of benefits returned *α* being the same for all host types (*α* = *f*(*α*) = *α_g_*), but *γ* > *γ_g_* = 1 > *h*(*γ*) such that a symbiont strain’s effectiveness in returning host investment is greatest when matched to the host specializing on it. Generalists have the same efficiency with both symbiont types. Thus, under cooperative specialization, more benefit is produced, but the relative division of it remains the same, such that both the specialist and its preferred symbiont benefit relative to the generalist and specialist with non-preffered symbiont.

In the antagonistic case, in contrast, specialist hosts have a higher *α* when paired with their preferred symbiont, specifically, 1 > *α* > *α_g_* > *f*(*α*) > 0. We assume in this case all pairings have the same *γ* (*γ* = *h*(*γ*) = *γ_g_*), such that there is no additional benefit being produced from specialization; specialist hosts are simply better at extracting a higher portion of the benefits from their preferred symbionts. This leaves symbionts that associate with matched specialist hosts with a lower fitness compared to interacting with generalist hosts or mismatched specialist hosts. We consider the term *γ* to be monotypic, meaning that there is no difference between phenotypes in the value of the benefit produced from symbiosis.

### 2.2 Spatial model

To understand the impact of spatial dynamics and heterogeneity, we extend these models by considering change in frequency of each type while diffusing across space. We model the diffusion process across the *xy*-plane by the following set of reaction-diffusion equations:

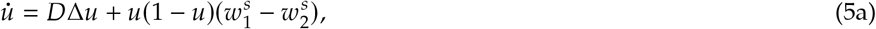

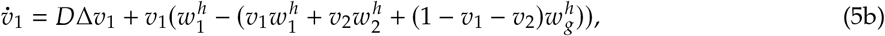

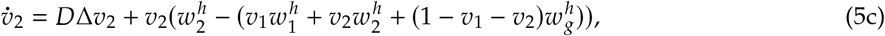

where *D* is the diffusion constant and Δ = *∂_xx_* + *∂_yy_* is the Laplace operator. See our Mathematica notebooks for the details of the numerical methods.

### 2.3 Invasion Analysis

Finally, we consider how the traits *α, γ*, and *x* will evolve using adaptive dynamics and numerical simulations. We examined the outcome of invasion of a rare mutant whose trait values (i.e. values of *α, γ*, or *x*) differed from those of residents (Brännström et al., 2013). Trait values favored by natural selection were those that allowed an initially rare mutant to increase in frequency. When specialization was cooperative, we determined the invasion exponent, the fitness of these rare invading mutant hosts or symbionts strategist relative to a resident population. We assumed that the changes in strategist frequency from our dynamical equations (4a)-(4c) and spatial model (5c) (ecological dynamics) occur rapidly relative to parameter changes (evolutionary dynamics). Thus, resident populations are the stable populations that emerge from the ecological dynamics. Next, we found the selection gradients, the change in the invasion exponent with respect to a change in a trait value *γ, α*, or *x*. The sign and stability of these selection gradients determined the pattern of selection that acted upon trait values. For mathematical details see SI-1.3. When hosts specialized antagonistically, invasion exponents were intractable because there was no stable resident population. Therefore, we employed numerical simulations to determine whether a rare mutant host or symbiont would successfully invade. See the our Mathematica notebooks and SI-2.2 for details.

## 3 Results

### 3.1 Cooperative specialization

#### 3.1.1 Ecological dynamics

The cooperative specialization model has two stable equilibria with matched host-symbiont pairs, i.e. all hosts are specialists of one type and all symbionts are their preferred symbiont. The system also features a continuous line of generalist host equilibria, which excludes specialist hosts, and has an intermediate symbiont frequency. The frequency of specialist symbiont 1 along this line, *u*\*, is determined by the difference between matched host payoff (*π*) and mismatched payoff (*π*′):

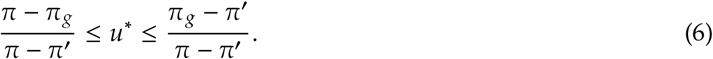

If the generalist payoff is less than the average specialist payoff (i.e. *π_g_* < (*π* + *π*′)/2), then this line of equilibria is unstable. If, however, generalist payoff is greater than the average specialist payoff, then it behaves as a type of saddle. It is attracting with respect to changes in the frequencies of hosts. However, symbiont frequencies may change by perturbations of the population. The endpoints of the line of equilibria are unstable. Thus, if the symbiont frequencies are perturbed beyond the endpoints, the system moves to one of the matching equilibria. Therefore, cooperative specialization is bistable in the long-run resulting in specialization in a well-mixed population (see SI-1.1 for the mathematical details and see Table SI-1 for relevant terminology). However, there can be long transients, depending on the frequency/intensity of mutations or invasions, during which all hosts are generalists.

In the spatially explicit model, on the other hand, we observe spatial polymorphisms. When hosts specialize cooperatively and spread across space from a distribution which is initially random, host and symbiont populations assort into monomorphic patches. The patches become stable over time when diffusion is sufficiently low as depicted in Figure 3. When diffusion is relatively high there is no stable patchy distribution, rather one specialist host and their preferred symbiont dominate the entire space (not depicted here). Which host-symbiont pair comes to dominate is determined by the initial abundances of all types.

**Figure 3.**
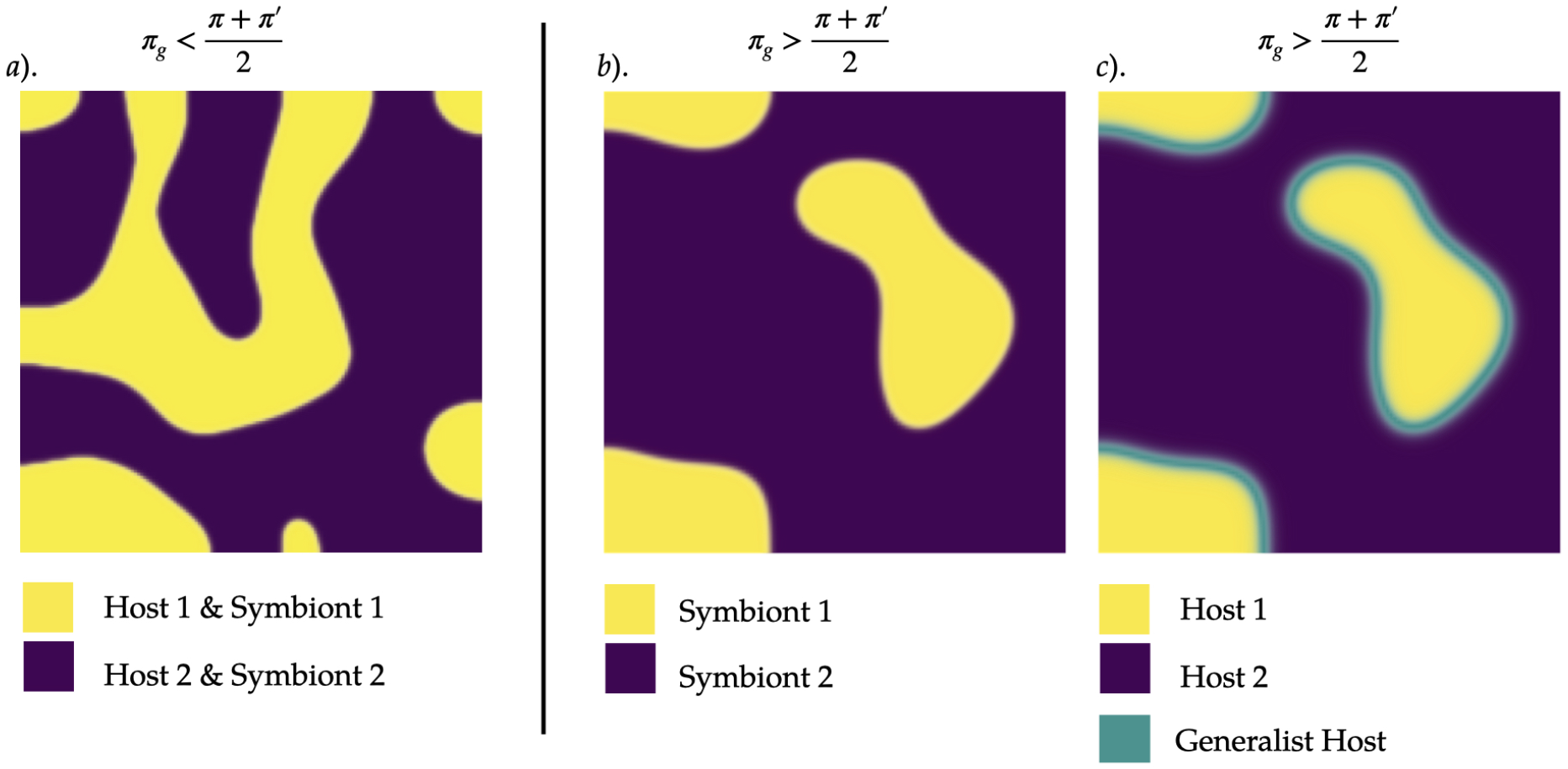
**When diffusion is low cooperative specialization leads matched symbiont-host pairings to fix in distinct patches across space. This figure depicts the results numerical simulations using the system of partial differential equations in (5c). In panel** *α* **specialist hosts exclude generalist hosts because their average payoff exceeds that of the generalist hosts. In panels *b* and *c* generalists may persist at borders between matched specialist patches. When generalist payoff exceeds the average payoff of a specialist, regions of intermediate symbiont frequency persist at the borders between matching host-symbiont patches (*b*). Generalist hosts are dominant at these intermediate symbiont frequency borders (*c*). In** *α*, *b*, **and** *c D* = 2.0 × 10^-6^, *α* = 0.5, **and** *x* = 0.5. **In panel** *a γ* = 0.1, *h*(*γ*) = 0.4, **and** *γ_g_* = 0.6. **In panels *b*) and emphc)** *γ* = 1.1, *h*(*γ*) = 0.1, **and** *γ_g_* = 1.

The presence of generalists in the spatial case depends on the condition *π_g_* > (*π* + *π*′)/2. When this condition is false, as depicted in of Figure 3-*a*), generalists are excluded. However, when the condition is true and diffusion remains low, generalists can survive at the border of patches as depicted in Figure 3-*c*. This phenomenon occurs because generalists are stable when symbiont frequencies lie within an intermediate range (given by the inequalities (6)), which is found at the boundaries between patches of different specialists. Increasing diffusion rates reduces the size of these boundaries, decreasing generalist frequency. However, if only symbiont diffusion increases relative to host diffusion, specialist hosts do not diffuse into borders as quickly as their preferred symbionts and thus the width of these regions is maintained. Furthermore, as the mix of symbiont types homogenizes, hosts are more likely to encounter the intermediate range of symbiont frequencies given the inequalities (6), which favors the generalist equilibria over the specialist equilibria. Because of this, generalists become more frequent as symbiont diffusion increases as depicted in Figure SI-1.

#### 3.1.2 Invasion analysis

As shown above, in a well-mixed population, the long-term ecological equilibrium under cooperative specialization is the fixation of one type of specialist host and their preferred symbiont. In the spatial model, too, each specialist and their corresponding symbiont is fixed locally, except for boundary regions. We first consider the evolution of symbiotic traits under this monomorphic scenario and then consider the boundaries in the spatial model where both symbionts might be present. The mathematical details of the analyses are given in SI-1.3.

To begin with, we assume that the symbiotic traits are under host control (i.e., evolve according to their effects on host fitness and not on symbiont fitness). In the monomorphic case we find that selection gradients for resource extraction *α* and efficiency *γ* are both positive, which means that mutants with higher *α* or *γ* can invade. This result is intuitive since *γ* increases the fitness of both the hosts and symbionts. And, because the symbiont population is monomorphic, the hosts do not experience the trade-off from increasing *γ* at the expense of *h*(*γ*). A higher *α* will similarly increase the amount of good provisioned for the host, and thus is also selected for in hosts. Finally, the host resource provision *x* evolves to the evolutionarily stable intermediate value of *αγ*/2, reflecting the balance of investment into symbionts versus the cost of this investment to hosts.

Next, for the uniform population comprised of a specialist and its preferred symbiont, we consider the parameters under symbiont control so that the fitness of invading symbiont mutants determines their evolution. The selection gradient for mutual efficiency, *γ*, and resource allocation to trade, *x*, are both positive, indicating that mutant symbionts with larger values of these traits can successfully invade and replace residents with lower values. However, the selection gradient for resource extraction, α, is negative, indicating that mutants with smaller values are favored by selection. This result is not surprising, as a symbiont is better off the more of the resource it keeps to itself. Further, this will have no indirect costs, since the host population is monomorphic and matching.

Our spatial ecological dynamics lead to monomorphic, matching host-symbiont patches. These patches have borders in which there is a mix of both symbiont types (Figure 3-*a*). Here, we consider how symbiotic traits may evolve in these border regions, where there is a mix of symbiont types. For analytic adaptive dynamics we can assume that the frequency of the non-evolving species is constant if the size of matching patches is large relative to borders, and diffusion of the non-evolving species is large relative to the mutating one. In this scenario, the patches act as a source of the non-evolving species at the borders: the frequencies of the non-evolving species at the border are determined primarily by diffusion from monomorphic patches. Thus, their frequencies will not change at the borders as the evolving border resident population evolves. First, we find that as in the monomorphic case larger values of *α* will evolve and *x* will evolve to an intermediate value of *αγ*/2. However, the resource efficiency *γ* is subject to a trade-off between matching and mismatching such that if *γ* increases (a host becomes more efficient with the preferred symbiont), *h*(*γ*) decreases (the host becomes less efficient with the non-preferred symbiont). We considered a variety of trade-offs (Supplemental Figure SI-2) to conduct a partial analysis for this case that assumes a fixed intermediate frequency of each specialist (see details in SI-1.3). We find that a concave up trade-off that increases at an increasing rate leads to invasion of ever larger values of *γ*. A concave down trade-off that increases at a decreasing rate, on the other hand, leads *γ* to converge to a stable value at which *h*′(*γ*) = *u*(1 – *u*). This concave down trade-off between matched and mismatched efficiency leads specialists to evolve to become a novel variety of generalist that receives approximately equal marginal benefit from interaction with either symbiont at the borders of fixed host patches. Finally, when *h*(*γ*) is linear, any value of *γ* may invade. These evolutionary analyses hold regardless of whether or not the evolving trait is under selection by the hosts or symbionts. Mathematical details are included in SI-1.3.

### 3.2 Antagonistic specialization

#### 3.2.1 Ecological dynamics

The antagonistic specialization model has two potential dynamical regimes. First, when generalist payoff exceeds average specialist payoff (i.e. *π_g_* > (*π* + *π*′)/2), specialist hosts are excluded and only generalist hosts remain at equilibrium. And, symbiont frequencies within the interval (6) form a stable line of equilibria. Within this interval, symbiont frequencies may drift from one value to another due to invasions or mutations. Conversely, when generalist payoff is less than the average specialist payoff (i.e. *π_g_* < (*π* + *π*′)/2), there are no stable equilibria, but a center at 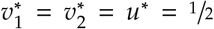. The frequencies of species oscillate around this intermediate point. The oscillations do not include the generalist hosts, as generalist hosts are always less fit than at least one specialist host. For the mathematical details of these results see SI-2.1.

Moving on to the spatial case, when *π_g_* > (*π* + *π*′)/2, generalists fix across space and both symbiont types coexist in each patch of the landscape, as symbiont diffusion leads their frequencies to homogenize across space after generalists fix. When *π_g_* < (*π* + *π*′)/2, however, there is no stable distribution of hosts and symbionts throughout space. Rather, the size and position of types of hosts and symbionts oscillate over time. Cycling occurs within each individual patch, and hosts and symbionts of each type diffuse from each patch at rates proportion to their frequency and the diffusion rate *D.* Gradually, the oscillations synchronize in each patch across space. We depict this result in Figure 4.

**Figure 4.**
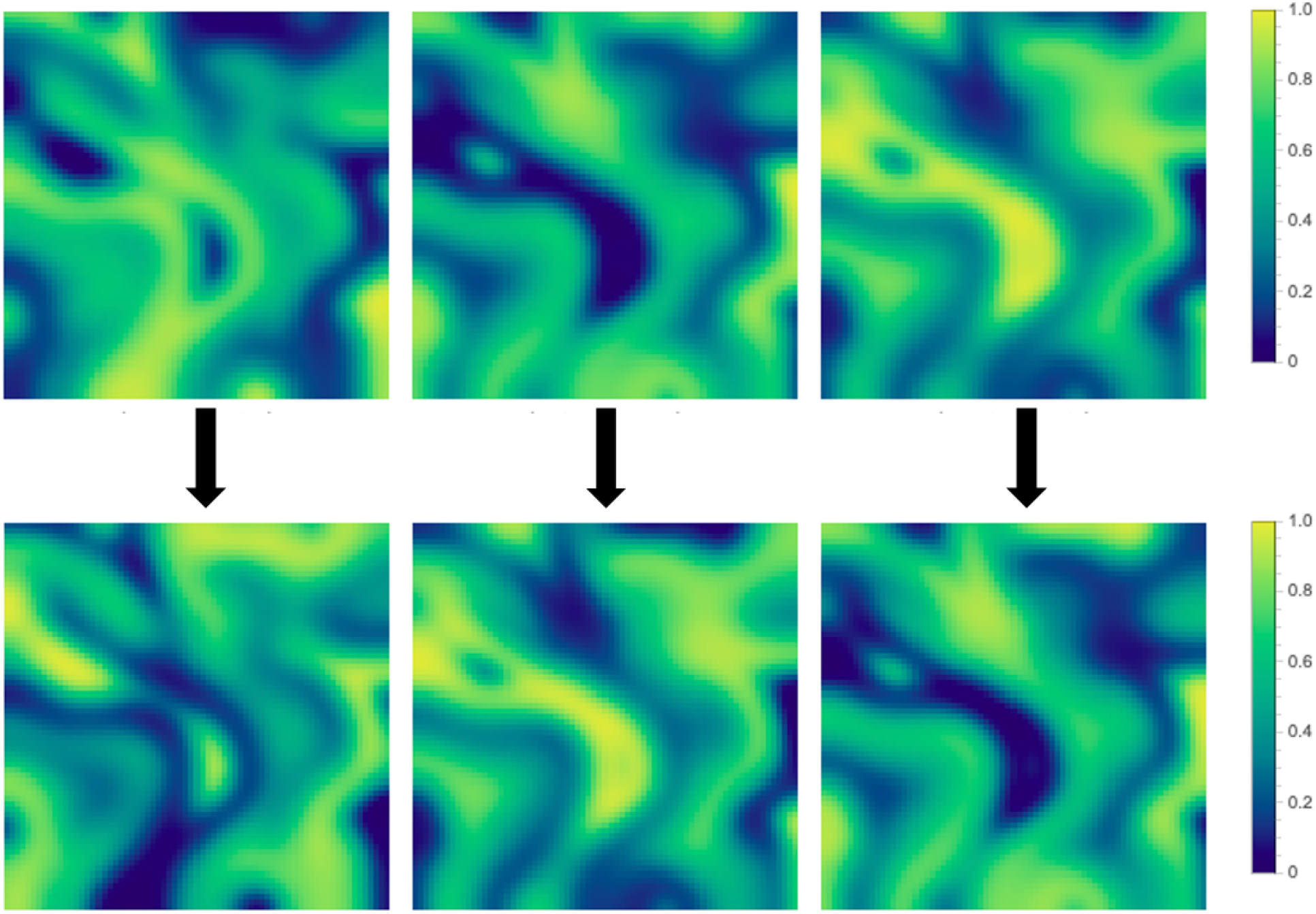
**When specialization is antagonistic, hosts and symbionts cycle from low to high frequency across space. Frequency of symbiont 1, host 1, and host 2 are given from left to right. The population begins with the spatial distribution of phenotype frequencies in the top row at an arbitrary time** (*t* = 672) **and competing types eventually come to occupy the spaces occupied by their competitors at a later point in the cycle** (*t* = 696). **Here** *D* = 10 *fα*) = 0.3, *α_g_* = 0.5, **and** *x* = 0.5.

The size of synchronous patches depends upon the value of the diffusion rate, *D*. When this rate is high, the entire plane cycles synchronously and its behavior matches the nonspatial case. When this rate is low, we observe patches cycling out of sync with one another. The lower the diffusion rate, the more distinct these patches become. When the diffusion rate differs between symbionts and hosts, the patch dynamics typically follow the partner with the higher diffusion rate. Greater symbiont mixing eventually results in greater host mixing, even when host diffusion is lower than that of symbionts. Similarly, host frequencies influence symbiont frequencies, thus greater host diffusion similarly results in synchronization with the greatest diffusion rate.

#### 3.2.2 Invasion analysis

For antagonistic specialization, our long term ecological dynamics in a well mixed population lead to a cycling population or to the generalist equilibria. First, we consider how the symbiotic traits (i.e. *α, γ* and *x*) will evolve in the cycling case. Analytic adaptive dynamics are intractable in this cycling population because the timing of mutant invasion dictates the frequency of the strategists with which mutants compete, and thus influences the success of invasion. We therefore used numerical simulations to consider mutations that may occur anytime along the cycle for our invasion analyses. Additional details describing these methods are included in SI-2.2.

In our first numerical simulations for the cycling population, we assume that *α, γ*, and *x* are under host selection and thus evolve according to their effect on host fitness. Because there is a mix of cycling symbionts, hosts experience a trade-off as mutations that increase the exchange rate (*α*) in matched symbiosis decrease *f*(*α*) in mismatched symbiosis. We considered linear, concave down, and concave up trade-off functions. For linear trade-offs, any value of *α* can invade a similar proportion of the cycle of a resident symbiont, indicating that α will evolve neutrally in this case (figure not shown). When *f*(*α*) is concave down there is a singular strategy 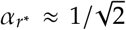 that cannot be invaded by any other nearby value of *α*. Thus *α_r_*\* is an evolutionarily and convergence stable strategy (Figure 5-*a*). In essence, hosts at *α_r_*\* are able to maximize average benefit from interactions with either symbiont type over the period of a cycle. On the other hand, when the trade-off is concave up greater values of α will be always favored for hosts (Figure 5-*b*).

**Figure 5.**
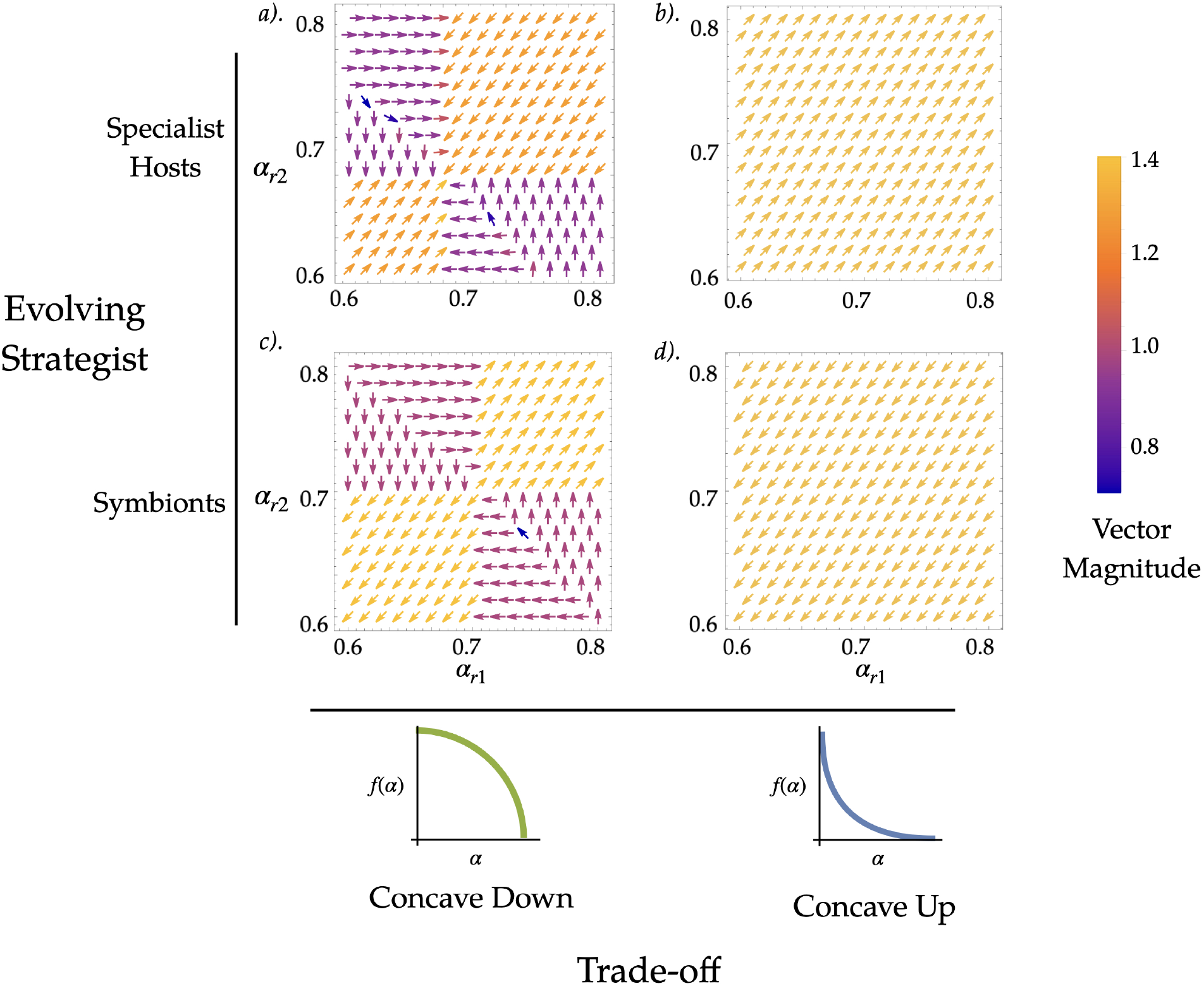
**In the cycling population, the outcome of mutant invasion is determined by the trade-off between matched and mismatched antagonism. The bottom row depicts the form of the trade-off functions we employed. Because both type 1 and 2 hosts and symbionts are present in the cycle, we considered mutants of both types. Panels *a-d* show vector plots whose horizontal component is the proportion of a cycle an invading type 1 mutant is able to invade and whose vertical component is the proportion of a cycle an invading type 1 mutant is able to invade. The plots in the top row are for initially rare invading specialist hosts, and plots in the bottom row are for initially rare invading symbionts. Here,** *x* = 0.5, **and** *γ* = 1. **The trade-offs are:** 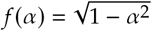 **(concave down), and** 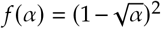 **(concave up). We used two sets of initial conditions:** *v*_1_ = 0.34, *v*_2_ = 0.66, *v*_1_ = 0.66, **and** *v*_1_ = 0.66, *v*_2_ = 0.34, *v*_1_ = 0.3

With respect to evolving *γ*, greater values of cooperation allow mutants to invade the cycle of a resident, thus higher values will always be favored. Finally, for the evolution of the proportion of traded resource (*x*), there is an evolutionary stable value of investment *x_r*_* ≈ (*α* + *f*(*α*))*γ*/4 that balances the costs and benefits from trade with both matched and mismatched symbionts. See SI-2.2 for the pairwise invasibility plots.

Next, we assume that symbiotic traits (i.e. *α, γ*, and *x*) are under symbiont selection in a cycling population. As with hosts, linear trade-offs between *α* and *f*(*α*) lead to neutral evolution of *α*. When the trade-off is concave down, there is an unstable singular strategy 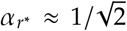 that can be invaded by nearby mutants (Figure 5-*c*). This leads the symbiont *α* to evolve away from the value of extraction that maximizes the fitness of the antagonistic hosts. A concave up trade-off will allow symbionts with reduced values of *α* to always invade the majority of a resident’s cycle (Figure 5-*d*). Essentially, selection pressure is on symbionts to escape the extraction of the host that specializes on them, at the expense of becoming more vulnerable to extraction from the opposite host type. Symbiont mutants with greater values of *x* and *γ* will always invade the majority of the cycle of a resident population. Greater resource investment (*x*) from hosts comes at no cost to symbionts, and there is no trade-off for increasing *γ* in the antagonistically specialized population. See SI-2.2 for the pairwise invasibility plots.

Our ecological dynamics also result in a community that is composed entirely of generalist hosts, and a mix of symbiont types when *π_g_* > (*π* + *π*′)/2. Because hosts and symbionts do not cycle in this scenario, we were able to determine selection gradients for symbiotic traits (*α, γ*, and *x*). In fact, because generalists traits do not vary for either symbiont type, the selection gradients for all symbiotic traits match those of the monomorphic case in 3.1.2. For mathematical details see SI-2.2.1.

## 4 Discussion

Our model provides predictions about antagonistic and cooperative specialization that have important applications to biological systems. Cooperative specialization tends to lead to host and symbionts specialized to one another. Though, generalists may prosper in the evolutionary short run or at spatial boundaries. Antagonistic specialization can lead to either generalist hosts or specialist hosts fixing, depending on the difference between the specialist host average payoff vs. the generalist host average payoff. Additionally, we considered the evolution of cooperative and antagonistic specialization. We found that increased cooperative specialization with a preferred partner evolves under cooperative specialization, while the evolutionary trajectory of antagonistic specialization depends on the trade-off between matched and mismatched antagonism in a cycling population.

When specialization is cooperative, our model predicts that host-symbiont co-diversification will occur. This is because under cooperative specialization, stable patches form in which only one specialist type is present (see Figure 3). The interior of these patches are isolated from gene flow from the conspecific specialist, thus there is reproductive isolation between strategists within each patch. Matched specialist partners will co-evolve to increase mutual benefit with their preferred partner at the expense of reduced benefit from mismatched symbiosis. This co-evolution will further reduce the viability of mismatched specialists that enter from other patches. This evolution further isolates conspecific specialist strategists and promotes pairwise speciation of the fixed hosts and symbiont in each patch. This dynamic will result in codiversified lineages of hosts and symbiont (Weiblen et al., 2015). By showing that cooperative specialization in mutualistic interactions may facilitate diversification, our model provides an important conceptual link between biodiversity and mutualism (Bascompte, 2019).

At the same time, specialization can also evolve to be antagonistic, with hosts or symbionts evolving to extract (or resist extraction of) more resources from each other. Such antagonistic specialization leads to cycling ecological dynamics in both well-mixed and spatially structured populations (Figure 4). The behavior of these cycles is determined by the trade-off between matched and mismatched antagonistic specialization. When this trade-off is concave up, natural selection favors increased resource extraction from symbionts, increasing the amplitude of the population’s cycle. Greater amplitude will eventually lead to the extinction of a specialist host or symbiont at cycle peaks or troughs. Alternatively, in the case in which generalists initially out compete specialists, it is possible for specialists to invade the generalist population if natural selection leads specialist payoff average to exceed generalist payoff. Thus, in the antagonistic model, specialization allows for consistent turnover of strategists across both time and space.

Our model is especially applicable to coral symbiosis, and provides predictions that are particularly relevant to two aspects of this interaction: 1) the degree of cooperation vs. antagonism exhibited in coral symbiosis; and 2) the conditions that favor specialization and generalism. Researchers have found evidence for cooperative and antagonistic interaction between corals and their symbiotic algae (Xiang et al., 2020) without clear resolution as to which pattern is more prevalent at the population scale. Understanding the relationship between these partners is especially important as mass mortality events (bleaching events) caused by anthropogenic climate change occur in scleractinian corals when the costs of symbiosis outweigh associated host benefits (Baker et al., 2018). Our model incorporates interactions at the scale of individual hosts and symbionts and predicts distinct population dynamics associated with both the cooperative and antagonistic interpretations of coral symbiosis. It is possible that further empirical observations of coral population dynamics with our predictions in mind may shed light on the prevalence of each mode of association in this symbiosis. The prevalence of generalism vs. specialism is also a matter of contention in cnidarian symbiosis. While most corals appear to be symbiont specialists (Poland and Coffroth, 2016), much evidence suggests that associations with other symbiont types are also viable (Silverstein et al., 2012). Our results that favor generalist hosts, which evolved under both antagonistic and cooperative specialization, indicate that coexistence of these two strategists amongst coral hosts is quite possible. In the cooperative case, generalists persist at the borders between specialists even in the absence of environmental heterogeneity. Generalism is also stable in the antagonistic case when average generalist benefit exceeds that of specialist benefit. The relative prevalence of these strategies also has important implications for the conservation of corals and other symbiotic species. Heat tolerance in corals is often linked to algal partner species identity, thus increased flexibility of host association may allow hosts to associate with algal partners with more robust cellular physiology (Berkelmans and Van Oppen, 2006). Alternatively, specific partners may be more likely to have physiology that is optimized for symbiosis with a preferred partner (Matthews et al., 2017). These partnerships may be better able to mutually co-evolve to more effectively resist thermally stressful events.

Our model can also be applied to the symbiosis between plants and mycorrhizal fungi to explain the prevalence of generalism in this particular partnership. Increased abundance of fungal partner species has been found to increase overall plant community diversity (Van der Heijden et al., 1998), and researchers have found evidence of host-symbiont specificity in this mutualism (Hoeksema et al., 2009), though generalism seems more prevalent. One reason for the prevalence of generalism could be the limited dispersal capability of plant and fungal partners. In our model, when dispersal rate is low (i.e. the diffusion constant D is low) ecosystems assort into a greater number of fixed host-symbiont patches, increasing the area of border regions in which an intermediate frequency of both symbiont partners occurs. In these border regions, generalism can evolve from specialism when specialization is cooperative. This result may explain why host and symbiont partners with large dispersive abilities such as corals and their zooxanthellic algae are often specialized (Poland and Coffroth, 2016), while plant-mycorrhizal partners tend toward generalism (Peay et al., 2015).

In our model, generalism persists in both antagonistic and cooperatively specialized mutualisms, indicating that this strategy may be adaptive even in the absence of ecological perturbation. Under antagonistic specialization, when generalist payoff is greater than the average specialist payoff, all hosts become generalists. In the case of cooperative specialization, mutually beneficial symbiosis means that increasing frequency of a specialist host simultaneously bolsters its preferred symbiont’s frequency. Thus, generalism, outside of spatial conditions, is unstable in the long run due to invasions and mutations. However, in spatial simulations, generalists persist at the borders between monomorphic specialist patches when generalist payoff is greater than average specialist payoff (Figure 3). Furthermore, when the evolution of continuous host trait values are considered, it is possible for generalism to evolve from specialization at these borders SI-2.2. These results show that generalists and specialists might co-coexist in ecological landscapes based on biotic interactions alone. In plant-microbial and plant-fungal mutualisms there is evidence that negative plant soil feedback promotes coexistence between a variety of plant, microbial, and fungal partner species, (Bever et al., 2010). This suggests precedent for coexistence between these strategies. In ecological communities, generalism could also be perpetuated by environmental disturbance. Events that reduce frequency of a preferred symbiont or host could create opportunities for generalist hosts to increase in frequency. In the case of mutualism, to be a generalist is to receive a benefit that is between the extremes that specialists receive from interacting with their preferred and not preferred partners. Generalists are therefore hedging their bets against the possibility that disturbance might make a preferred partner species unavailable. Interestingly, our model shows that generalism can be evolutionarily stable even in an undisturbed environment. Incorporating ecological feedback into later iterations of this model could provide interesting insights into the adaptive benefits of generalism.

In mutualistic interactions in which specificity and generalism have complex benefits and costs, it is difficult to determine how these strategies will evolve. Our model provides critical insights into how specialization and generalism will evolve in both cooperative and antagonistically specialized mutualisms involving the exchange of a good. Our results showing that these strategies can coexist in cooperative mutualism, and that the frequency of each strategist cycles in antagonistic mutualism, reveal how different modes of species interaction have distinct outcomes. Tying together the degree of specificity and cooperation in a species interaction has important implications for numerous mutualisms that are critical to both biodiversity (e.g. coral-algal mutualism) and agricultural productivity (e.g. legume-rhizobia mutualism). Further study examining the implications of our model may continue to enhance understanding of these systems.

## Conflicts of interest

The authors declare that they have no conflicts of interest.

## Acknowledgements

The authors would like to thank the Akçay, Barott, and Plotkin labs for helpful feedback and discussions.

## Author contributions

All authors contributed to the conception of the study and the final manuscript. C.C. developed the code for and analyzed the numerical simulations. C.C. and B.M. carried out the mathematical analysis and wrote the first draft.

## Financial support

This research was supported by the University of Pennsylvania.

## Data availability

The code for the stability analysis and numerical simulations were written in Mathematica and can be found at https://github.com/christopheriancarlson/SpecificMutualism.

## Supplementary Information

Here we detail the invasion and stability analyses for cooperative (SI-1) and antagonistic (SI-2) specialization. For the stability analysis, Table SI-1 holds a glossary of our terminology.

**Table SI-1.**
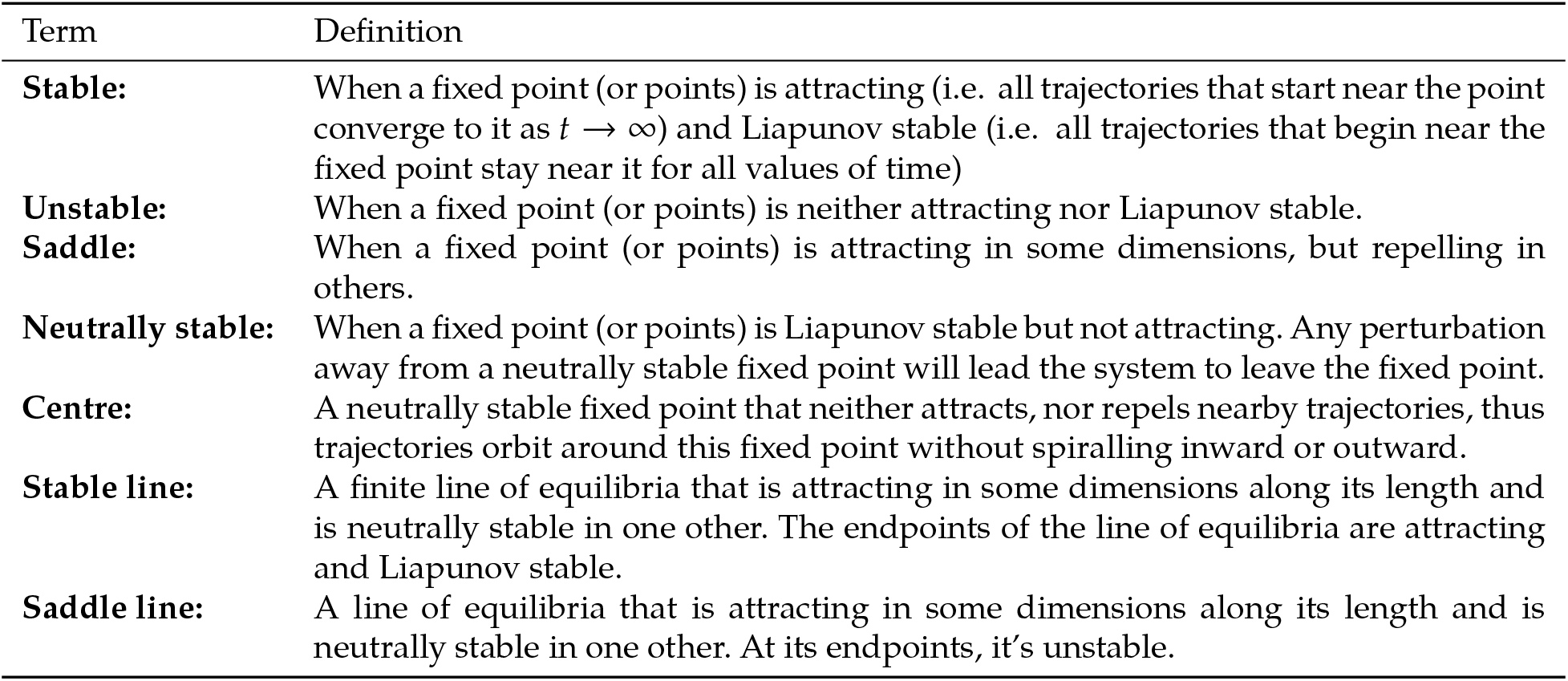
Glossary of stability analysis terminology.

### SI-1 Cooperative specialization

We begin with stability analysis of equilibria (SI-1.1), followed by simulations of the spatial reaction-diffusion system and its impact on generalist frequencies (SI-1.2), and evolutionary invasion analyses of mutations in the traits *α, γ*, and *x* (SI-1.3).

#### SI-1.1 Stability Analysis

Here we detail the stability results of the cooperative specialization model. The payoffs to specialist hosts for matched and mismatched pairings are *π* and *π*′, respectively. Generalist hosts receive the payoff *π_g_* regardless of which symbiont they are paired with. Symbiont payoffs are *ψ* and *ψ*′ for matched and mismatched pairings, respectively, and *ψ_g_* for when they are paired witht he generalist host. In cooperative specialization, *α* = *f*(*α*) = *α_g_* and *γ* > *γ_g_* > *h*(*γ*) > 0. We thus have the following payoffs:

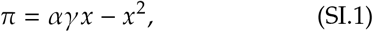

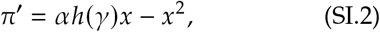

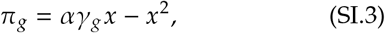

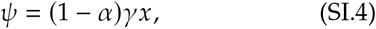

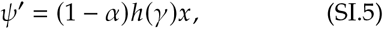

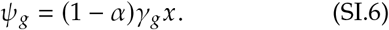

The average payoffs, 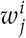 for *i* = *h, s* (host or symbiont) and *j* = 1,2, *g* (specialist 1 or 2, or the generalist host), are given by the payoffs and frequencies of specialist hosts (*v*_1_ and *v*_2_) and symbiont 1 (*u*). Payoffs are thus:

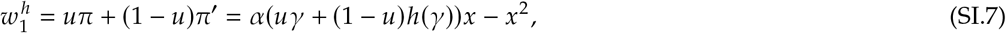

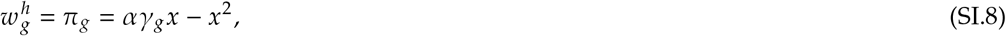

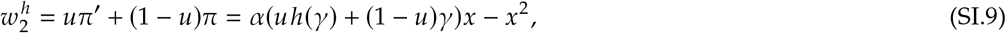

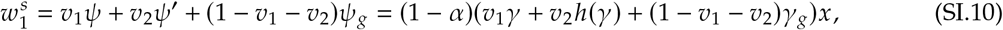

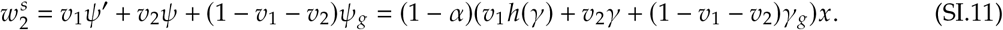

We may now write the complete equations for cooperative specialization:

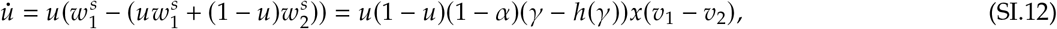

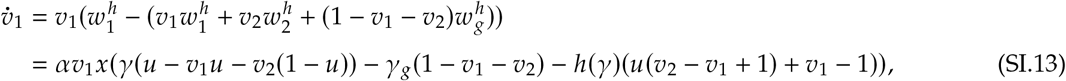

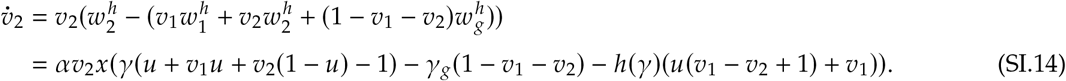

Note that equation SI.12 implies that the frequency of symbiont 1 will increases if there are more of host type 1 than host type 2, and otherwise decrease.

We linearize the system and evaluate the eigenvalues of the Jacobian at the equilibria. We first consider the boundary equilibria, i.e. equilibria with monomorphic host and monomorphic symbiont populations, followed by the polymorphic equilibrium. We denote equilibria using ordered triples: (*u*, *v*_1_, *v*_2_). We summarize the stability conditions for the equilibria in Table SI-2. In summary, cooperative specialization leads to bistability where the population is composed of monomorphic populations of hosts and symbionts who are matched to one another. However, there is the possibility of long transients of generalists with polymorphic population of symbionts. If *π_g_* > (*π* + *π*′)/2, there exists a set of equilibria that form a line with a mix of both types of symbionts and only generalist hosts. This line is attracting or neutrally stable along its length, but unstable at its endpoints. Thus, in the long term under perturbations, it is unstable.

**Table SI-2.** Summary of stability of equilibria for cooperative specialization. See Table SI-1 for a glossary of these terms.

| Equilibria | Stability |
| --- | --- |
| **Boundary equilibria** |  |
| Matched host and symbiont type: $(0, 0, 1), (1, 1, 0)$. | Always stable node. |
| Mismatched host and symbiont types: $(0, 1, 0), (1, 0, 1)$. | Always unstable node. |
| Generalist host and monomorphic symbionts: $(0, 0, 0), (1, 0, 0)$. | Always a saddle node. |
| **Interior (polymorphic) equilibria** |  |
| Generalist host and polymorphic symbionts: $(u^*, 0, 0)$. | If $\pi_g > (\pi + \pi')/2$, then a saddle line.<br>Else, unstable. |
| Coexisting specialist hosts and symbionts: $(1/2, 1/2, 1/2)$. | Always unstable. |

##### SI-1.1.1 Boundary equilibria

At (0,0,1) and (1,1,0) the specialist host and its preferred symbiont are fixed. These are matched equilibria and the corresponding eigenvalues are:

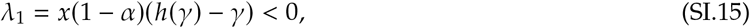

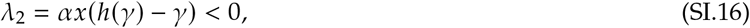

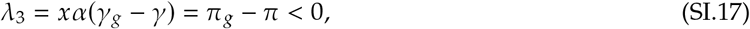

and thus these matched equilibria are stable. However, the mismatched cases, (0,1,0) and (1,0,1), are unstable, since the eigenvalues of the Jacobian are all positive:

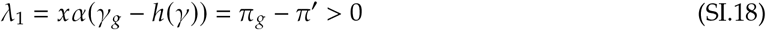

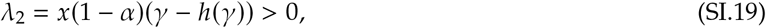

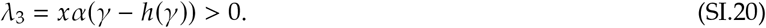

Consider next the equilibria (0,0,0) and (1,0,0), wherein our population is comprised of generalist hosts and only one symbiont type. The eigenvalues of the Jacobian matrix evaluated at either of these points are:

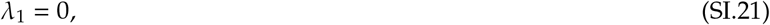

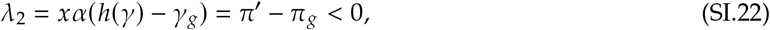

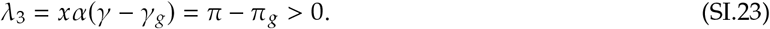

Therefore, these equilibria are unstable. Note that when 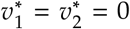, *u*\* may be any value in [0,1], which brings us to the internal equilibria.

##### SI-1.1.2 Internal equilibria

If 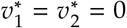, then we have a line of equilibria, *u*\* ∈ [0,1]. i.e. all hosts are generalists and symbionts may be any combination of each type. The corresponding eigenvalues for this line of equilibria are

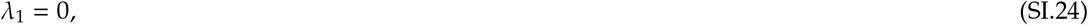

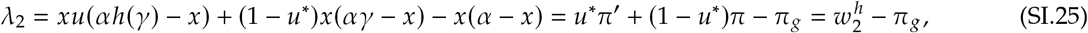

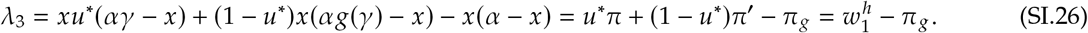

(SI.25) and (SI.26) can both be negative when

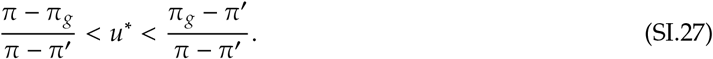

This inequality will be satisfied when *π_g_* > (*π* + *π*′)/2 resulting in an interval where both eigenvalues are negative.

Within the interval SI.27, the line of equilibria is attracting. Outside of it, it is repelling. We apply the theory of bifurcation without parameters to explore the effects at the end points (Liebscher, 2015). Without loss of generality, consider the right end point of SI.27. We can change coordinates such that bifurcation variable *μ* = *u* + (*π*_g_ - *π*′)/(*π* - *π*′), *y* = *v*_1_, and *z* = *v*_2_. We have equilibrium at (*μ,y,z*) = (0,0,0). The Jacobian is

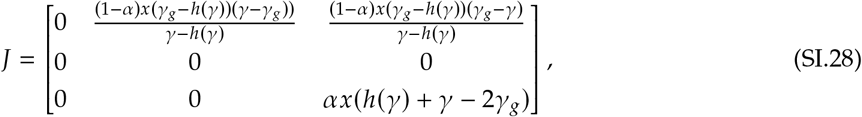

which is in the desired form.

Next we use the Center Manifold Theorem to characterize the stability of the higher order terms of the system as a function of the bifurcation parameter *μ* and the centre dimension *y*, i.e. *z* = *m*(*μ,y*) = *α*_11_*μy* + *a*_20_*μ*^2^ + *a*_02_*y*^2^ + .… To solve for the coefficients *a_ij_* we use the invariance condition

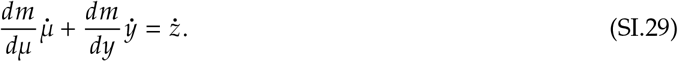

From here, however, we find that all coefficients must be zero and thus *m*(*μ, y*) = 0. The dynamics on the centre manifold are thus described by

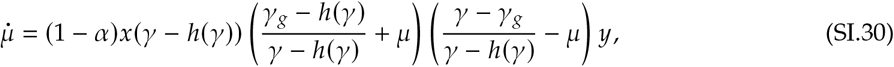

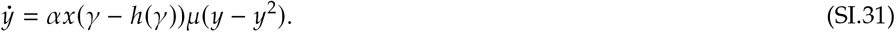

This system give us a transcritical bifurcation without parameters, i.e. the end points of the stable interval are semi-stable (there are nearby regions which are attracted to the line of equilibria and the bifurcation point and regions where we are repelled). The bifurcation point at the other end of the interval is also semi-stable.

We now consider internal equilibria 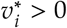. Equations (SI.13) and (SI.14) are at equilibrium when

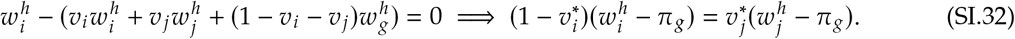

If 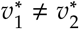, then Equation SI.12 is zero if *u*\* = 0 or *u*\* = 1. This state corresponds to monomorphic symbiont and polymorphic host populations. Without loss of generality, let *u* = 1, then 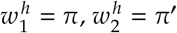, which gives us

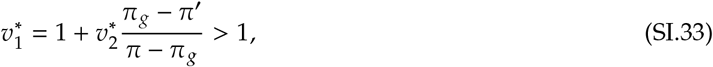

since 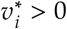. Thus, it is impossible for our system to have such an equilibrium.

Now consider equilibria where 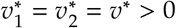. Equation SI.32 with *i* = 1 and *j* = 2 gives us

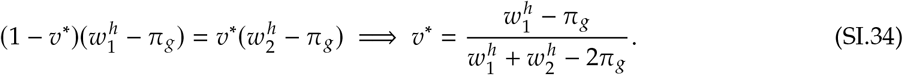

Then, substituting *v*\* into Equation SI.32 with *i* = 2 and *j* = 1, we have

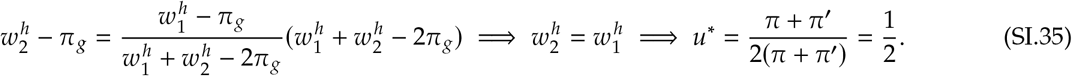

Then, 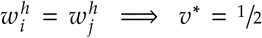. Our final polymorphic equilibrium is thus (1/2, 1/2, 1/2). The corresponding eigenvalues are:

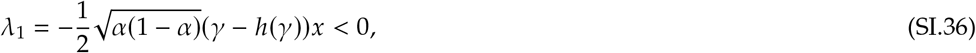

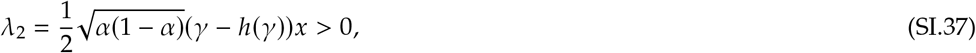

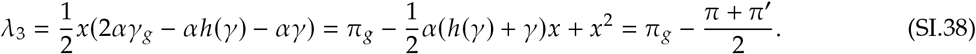

Though Λ_3_ may be either positive or negative depending on our assumptions, this state is a saddle since Λ_1_ < 0 and Λ_2_ > 0.

#### SI-1.2 Spatial simulations

We numerically simulated the spatial model from (5c) for cooperative specialization. These simulations are included in supplemental Mathematica notebooks found at https://github.com/christopheriancarlson/SpecificMutualism. Here, we detail the results of these simulations when generalist payoff is greater than average specialist payoff, i.e *π_g_* > (*π* + *π*′)/2. We examined what diffusion rates would allow generalists to persist when this inequality is satisfied. Recall that under this condition the generalist equilibrium is a saddle line, and when this condition isn’t met, generalists are completely excluded.

In all simulations, we varied the diffusion rate of strategists and measured the average frequency of generalists in an *xy* plane across which strategists were diffusing after letting our simulation run for a significant number of time steps (*n* = 3,000). We considered 20 randomly generated initial frequencies of hosts and symbionts across the plane, and calculated summary statistics for these 20 simulations. We systematically varied the diffusion rate for three cases. In the first, all types have the same diffusion rate and this rate is varied. In the second case, symbiont diffusion rates are fixed, and only host diffusion rates are varied (note that all hosts have the same diffusion rate). In the third case, host diffusion rates are fixed, and only symbiont diffusion rates are varied.

Figure SI-1 depicts our results. When varying the diffusion rate of all strategists or only hosts, generalist frequency decreases as the diffusion rate increases (panels a and b). This occurs because higher diffusion rates reduce the width of the borders between patches of matching hosts and symbionts. However, further increasing the diffusion rate results in a dramatic increase in generalist frequency. Since, the high degree of diffusion is causing patches to become polymorphic. As the strategists become more well mixed across space, generalists are favored.

When only symbiont diffusion increases, the frequency of generalists increases (Figure SI-1 c). Since hosts from monomorphic patches do not diffuse into borders as rapidly as their favored symbionts, the ecological dynamics at the borders do not drive them to being monomorphic. Thus, the width of the borders between patches does not decrease. However, the mix of symbiont types homogenizes, which favors the generalist equilibria over the specialist equilibria.

#### SI-1.3 Evolutionary invasion analysis

Parameters are analyzed where they are under selection by either hosts or symbionts; i.e. mutating traits are assumed to be a trait of either the host or symbiont and not both. We first consider evolution at the matched pairing equilibria and then at the saddle line where generalist are present. When *π_g_* > (*π* + *π*′)/2, the generalist equilibria is unstable. However, in the spatial scenario, the border between monomorphic matched patches can support generalists. We consider invasion analysis under this special case in SI-1.3.3. Table SI-3 summarizes the results.

**Figure SI-1.**
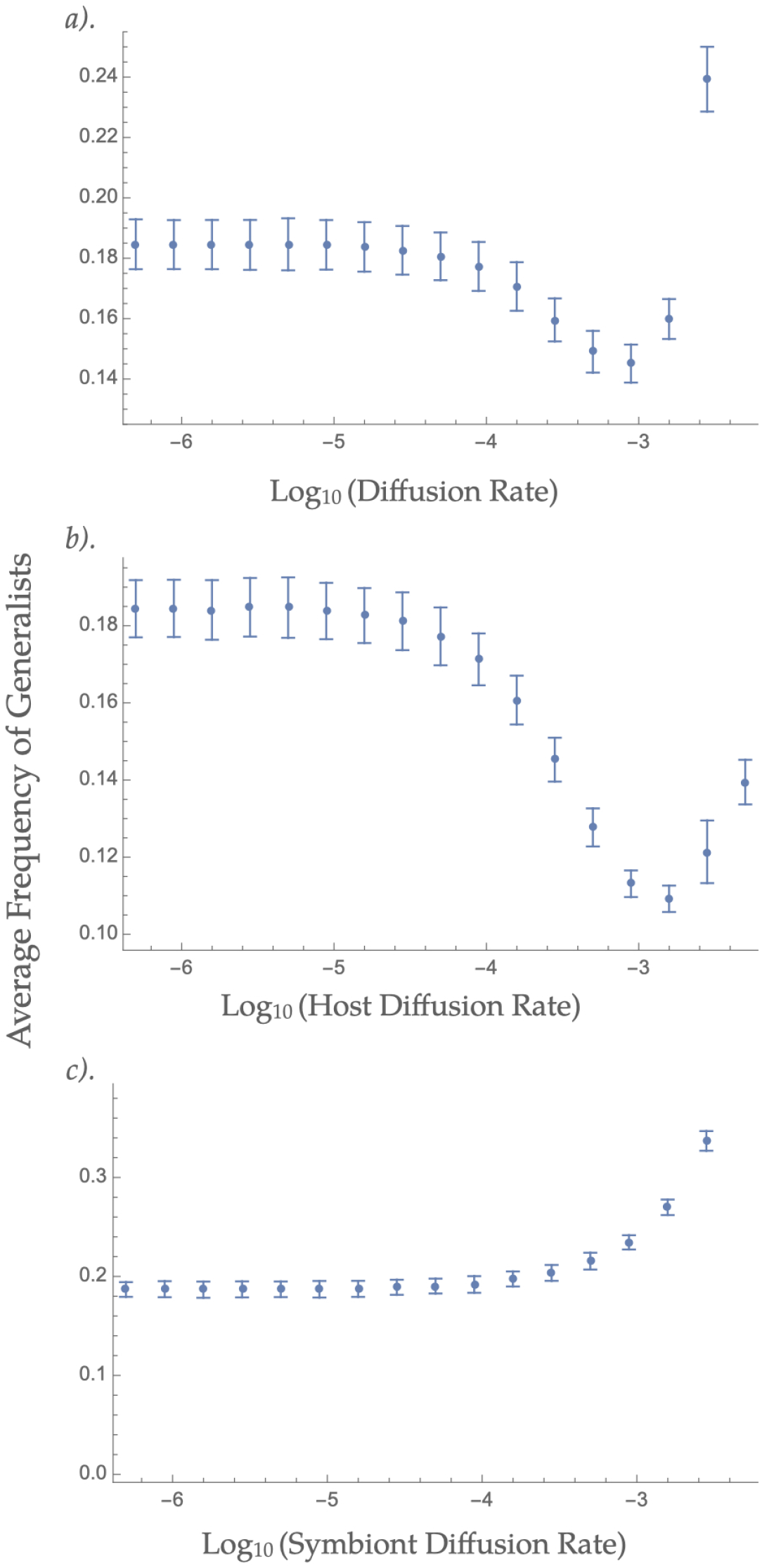
**The average frequency of generalist hosts across 20 simulations when the diffusion rate varied. Error bars denote the the standard error of the average across these. In *a*). the diffusion rate of all hosts and symbionts varied. In *b).* only the diffusion rate of hosts was varied, while symbionts all diffused at a constant rate (***D* = 5.0 × 10^-5^). **In** *c*). **the diffusion rate of symbionts was varied, with a constant host diffusion rate of** *D* = 5.0 × 10^-5^

**Table SI-3.** Summary of invasion analyses for cooperative specialization. Note that the invasion analysis at the monomorphic and generalist equilibria are the same (with the caveat that perturbations could knock the system away from the generalist equilibria).

| Parameter | Host selection | Symbiont selection |
| --- | --- | --- |
| $\alpha$ | Greater values invade. | Lesser values invade. |
| $\gamma$ | Greater values invade. | Greater values invade. |
| $x$ | ESS and CSS at $x_{r^*} = \alpha_r \gamma_r / 2$. | Greater values invade. |

| Parameter | Trade-off | Host selection | Symbiont selection |
| --- | --- | --- | --- |
| $\alpha$ | n/a | Greater values invade. | Lesser values invade. |
| $\gamma$ | concave down | ESS and CSS at $h'(\gamma_{r^*}) = -u(1 - u)$. | ESS/CSS at $h'(\gamma_{r^*}) = -u(1 - u)$. |
| $\gamma$ | linear | Evolve neutrally. | Evolve neutrally. |
| $\gamma$ | concave up | Greater values can invade. | Greater values can invade. |
| $x$ | n/a | ESS and CSS at $x_{r^*} = \alpha_r \gamma_r / 2$. | Greater values invade. |

(b) Invasions on the spatial boundary between different monomorphic patches. ESS and CSS mean evolutionarily stable singular strategy and convergence stable singular strategy, respectively.

##### SI-1.3.1 Evolution at a matched pairing equilibrium

Under cooperative specialization, the long-term ecological dynamics in a well-mixed population result in one specialist host and its matching symbiont fixing. Here we consider whether a mutant can invade in such a case. We first consider the scenario where the trait is under selection by the host, and then by the symbiont. Thus they evolve according to their effect on host (symbiont) fitness. The invasion exponent, the growth rate of a rare mutant host (whose trait values are denoted with the subscript m) relative to the resident population that is composed entirely of a specialist host and its matching symbiont, is given by:

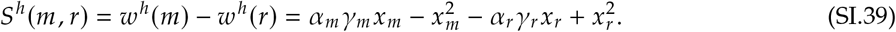

The subscripts *m* and *r* denote mutant and resident trait values, respectively. The *h* superscript denotes hosts. Here we avoid the use of a subscript for the host type, since the analysis holds whether we talk about one or the other. The selection gradients for *α_m_* and *γ_m_* are:

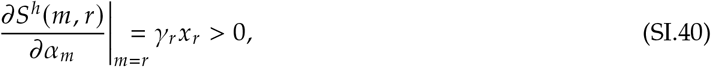

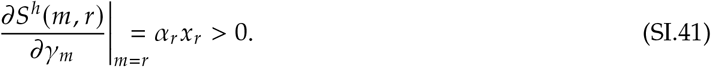

Because these selection gradients are positive, invading host mutants with larger values of *α* or *γ* than resident hosts will always be able to supplant native individuals. For *x_m_*, the selection gradient is

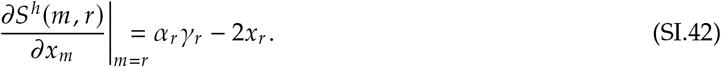

This is positive if 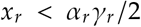, and negative when the inequality is flipped. 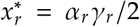 is a singular strategy (i.e. the selection gradient is zero). To check for evolutionary stability at this point and whether it is evolutionarily and convergence stable, respectively, we take:

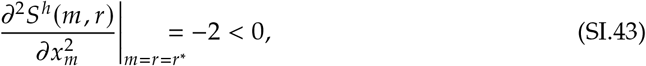

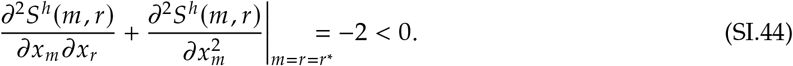

Since both are negative, 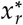 is both an evolutionarily and convergent stable state.

Now we assume that the symbiotic traits are under selection by the symbionts, and will evolve according to their effects on symbiont fitness. The invasion exponent for a symbiont that is fixed in a population together with its matched resident is

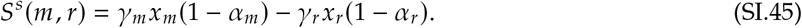

The *s* superscript denotes an invading symbiont. The selection gradients for *γ_m_, x_m_* and *γ_m_* are:

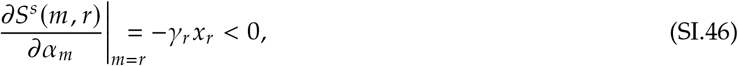

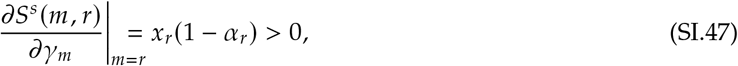

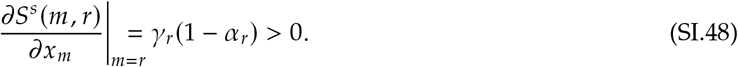

Positive selection gradients for *γ* and *x* indicate that selection will always favor fixation of symbiont mutants with greater values of these traits. Because the selection gradient for *α* is negative, mutants with reduced values will always invade and fix successfully.

##### SI-1.3.2 Evolution at generalist equilibria

Here, we consider how natural selection will act on host and symbiont trait values at the saddle line equilibrium. Note that this equilibrium is unstable at the endpoints of the line. Thus, mutations could push population to one of the matched pairing equilibria. Our results discussed here thus only hold so long as we remain within the line.

First, we consider the invasion exponent for an invading generalist host:

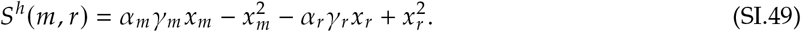

The *m* subscript denotes a mutant trait value and *h* denotes that this is the invasion exponent for hosts. Here, the evolving hosts are generalists that receive *γ_g_* from all symbionts; however, for this analysis subscripts will denote whether or not the value of *γ* belongs to a mutant or resident. Note also that since they receive the same payoff from both symbionts, *S^h^*(*m, r*) is not a function of *u*.

The selection gradients for *α_m_* and *γ_m_* are:

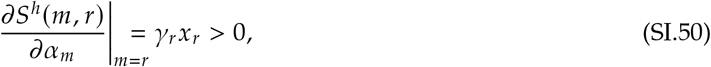

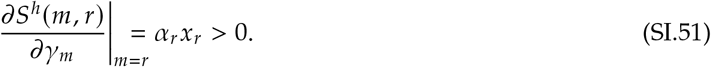

Note that the *r* subscript denotes the trait values of residents. Positive selection gradients indicate that invading host mutants with larger values of either *α* or *γ* will always successfully invade. Note that because the generalist receives the same *γ* from either symbiont type, the selection gradients for these traits do not depend on symbiont frequency or a match-mismatch trade-off. The *x_m_* selection gradient is

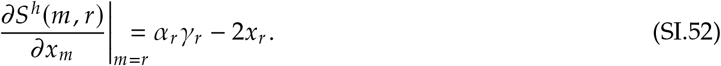

There is a singular strategy 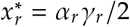. Checking for evolutionary and convergence stability we have

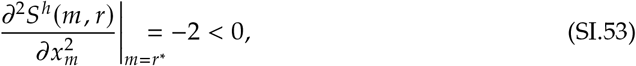

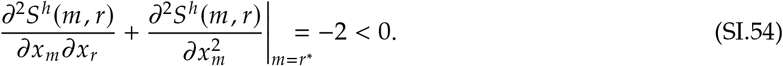

Thus, 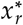 is a an evolutionarily stable local fitness maximum for generalist hosts that is convergence stable.

Next we consider how selection will act on the traits of symbionts at this generalist equilibrium. We consider first the invasion exponent for an invading symbiont:

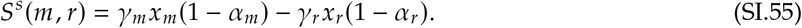

The superscript *s* denotes that this invasion exponent is that of a symbiont. The selection gradients are:

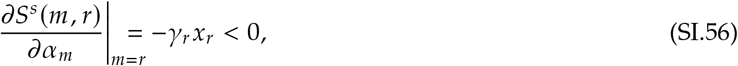

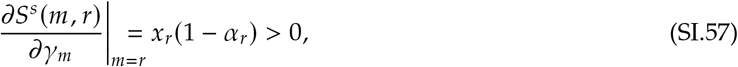

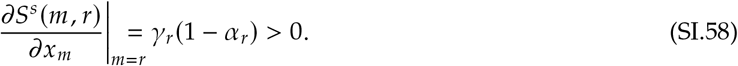

The negative selection gradient for α indicates that symbionts with lower levels of *α* are favored; they are better able to escape host extraction. The positive selection for *γ* and *x* indicate that symbionts with larger values of these traits will always be able to invade.

##### SI-1.3.3 Evolution at spatial boundaries

The spatially explicit ecological dynamics result in patches of specialist hosts and their matching symbionts. Within these patches, the monomorphic analyses as detailed above apply; specifically hosts experience no trade-off from specialization on their preferred symbionts, because the preferred symbiont is fixed. However, at the boundaries of these patches are regions where both symbionts might be maintained at positive frequency, and therefore hosts will experience the trade-offs associated with specialization. We now look at how selection may act in these regions. Specialists of different types coexist at these borders due to flow from the regions with matched symbionts and hosts. To analyze the invasion of mutant hosts/symbionts in these boundaries, we make the simplifying assumptions that the frequencies symbionts/hosts remain fixed. This assumption can be regarded as the case where the boundary is small relative to the size of and influx from the monomorphic patches that it fringes. Thus, the frequencies of the type not under invasion analysis will not change as the border resident population evolves. The invasion exponent for an invading host mutant at such a boundary is

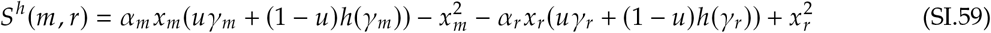

with 0 < *u* < 1 being fixed. This gradient does not consider the relationship between selection and the frequency of strategists across space, rather it assumes non-zero frequencies of specialist host and symbiont phenotypes; therefore, this is a partial analysis that only applies to borders. The selection gradient for the specialist host with respect to *α_m_* is

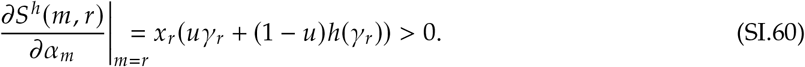

Because this selection gradient is positive, mutants with larger values of *α* will always successfully invade. The selection gradient for *γ_m_* is

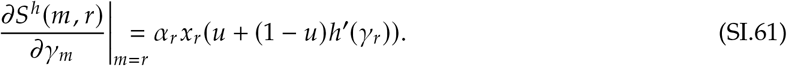

The sign of this function depends upon the trade-off function *h*(*γ*). Larger values of *γ* will be favored if *h*′(*γ*) > -*u*/(1 – *u*) and smaller values of *γ* will be favored if *h*′(*γ*) < -*u*/(1 – *u*). We considered concave up, linear and concave down trade-off functions as depicted in Figure SI-2, which depicts example trade-offs and their first and second derivatives.

If a value *γ_r*_*. exists at which *h*′(*γ_r*_*) = -*u*/(1 – *u*) then it is possible that *γ_r*_*. is a singular strategy. Checking for evolutionary and convergence stability, we have:

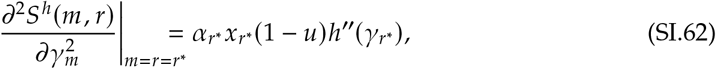

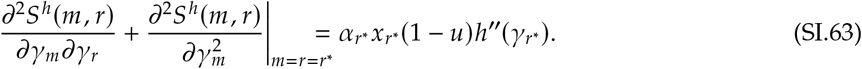

When the trade-off is linear *h*″(*γ_r*_*) = 0, thus *γ_r_* will evolve neutrally at the singular strategy. When the trade-off is concave up, *h*″(*γ_r*_*) > 0 and thus evolutionary branching will occur: the population will become dimorphic since this point is convergence stable. When the trade-off is concave down, *h*″(*γ*) < 0 and thus if there is a singular strategy, then it is both evolutionarily and convergence stable.

Next consider *x_m_*. The selection gradient is

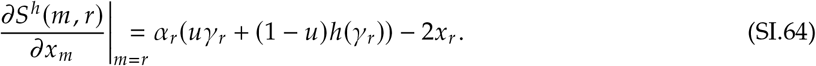

**Figure SI-2.**
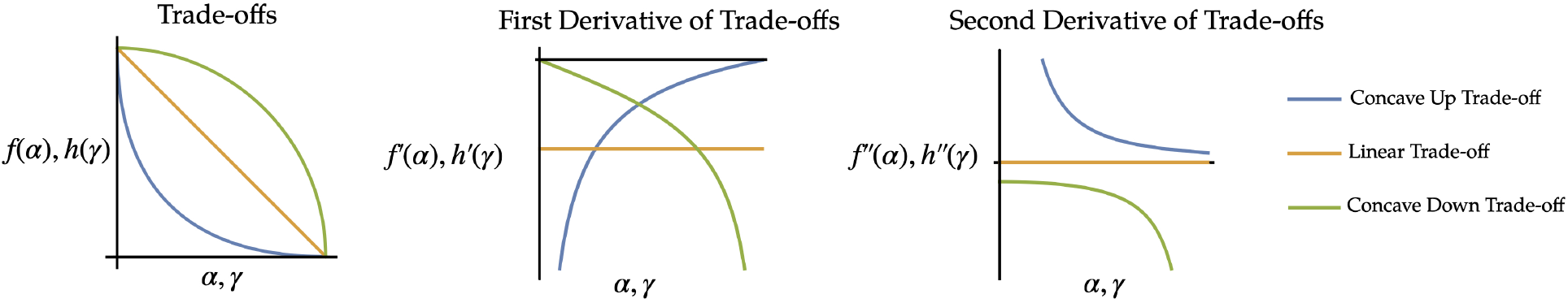
**Visualization of concave down (green), linear (yellow), and concave up (blue) trade-off functions considered for** *α* **and** *γ*. **The first derivatives of the trade-off functions determine the pattern of selection acting upon** *γ* **or** *α*, **while the second derivatives of the trade-off determines the evolutionary stability of values of** *γ* **or** *α*.

There is an evolutionarily and convergence stable singular strategy 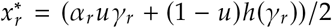. Since, evaluating the second derivative, we have

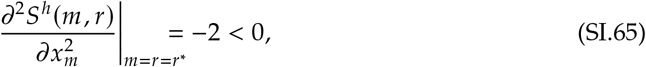

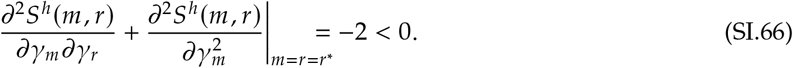

Now consider the fitness of an invading symbiont mutant at a border between specialist patches where symbionts of type *i* associate with matched host type *i* or mismatched type *j* i.e.

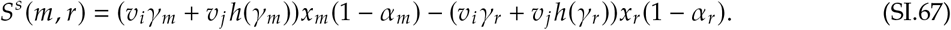

The selection gradients for *α, γ*, and *x* are:

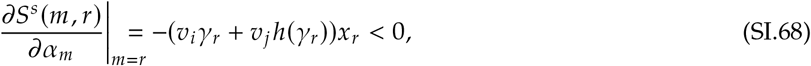

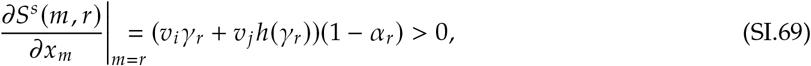

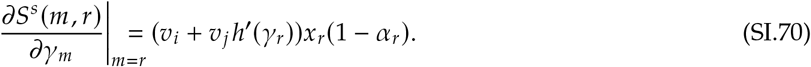

Selection will favor reduced values of α and increased values of *x*. The selection gradient for *γ* is positive or negative depending on which side of *h*′(*γ_w_*) = –*ν_i_/ν_j_* we are on. There is a singular strategy *γ_r_\** if *h*′(*γ_r_\**) = -*ν_i_/ν_j_*. We evaluated the evolutionary stability of this strategy by calculating the second derivative:

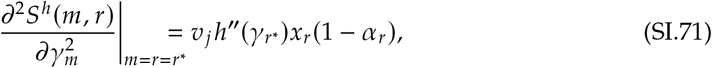

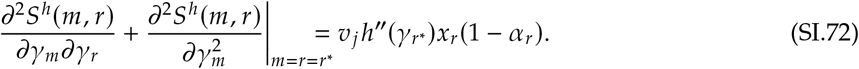

The trade-off *h*(*γ*) will determine the sign of the second derivative. When the trade-off function is linear, *h*″(*γ_r_\**) = 0 and *γ* will evolve neutrally. When the trade-off is concave up, *h*″(*γ_r_\**) > 0 and evolutionary branching will occur leading to *γ* dimorphism. Finally, when *h*″(*γ_r_\**) < 0 the singular strategy 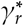 is both evolutionarily and convergence stable.

### SI-2 Antagonistic Specialization

We begin with stability analysis of equilibria (SI-2.1) followed by the invasion analysis (SI-2.2).

#### SI-2.1 Stability analysis

Here we detail the stability results of the antagonistic specialization model. The payoffs to specialist hosts for matched and mismatched pairings are *π* and *π*′, respectively. Generalist hosts receive the payoff *π_g_* regardless of which symbiont they are paired with. Symbiont payoffs are *ψ* and *ψ*′ for matched and mismatched pairings, respectively, and *ψ_g_* for when they are paired with the generalist host. In antagonistic specialization, *γ* = *h*(*γ*) = *γ_g_* and 1 > *α* > *α_g_* > *f*(*α*) > 0. We thus have the following payoffs:

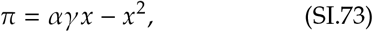

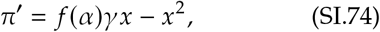

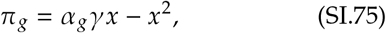

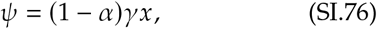

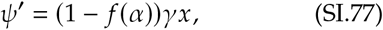

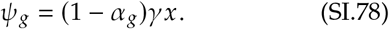

The average payoffs, 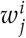 for *i* = *h, s* (host or symbiont) and *j* = 1,2, *g* (specialist 1 or 2, or the generalist host), are given by the payoffs and frequencies of specialist hosts (*v*_1_ and *v*_2_) and symbiont 1 (*u*). Payoffs are thus:

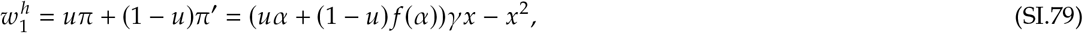

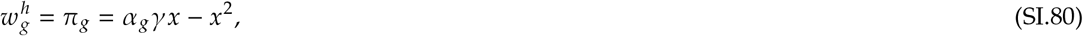

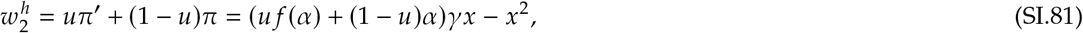

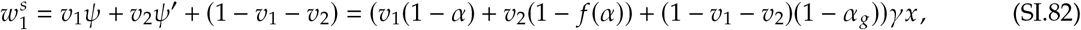

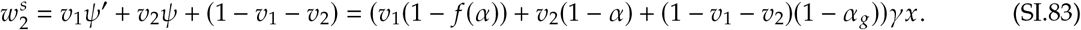

We may now write the complete equations for antagonistic specialization:

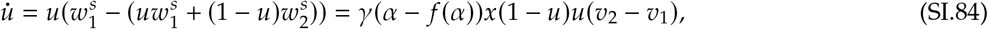

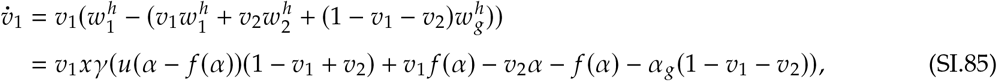

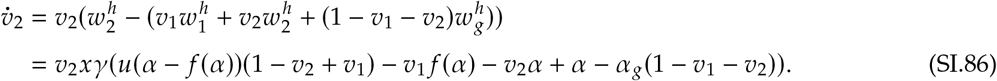

As in Section SI-1.1, we linearize the system and evaluate the eigenvalues of the Jacobian at the equilibria. We initially consider the boundary equilibria (monomorphic host and symbiont populations) followed by a polymorphic equilibrium. Equilibria are denoted using ordered triples: (*u*, *v*_1_, *v*_2_). We summarize the stability conditions for all of the equilibria in Table SI-4. In summary, antagonistic specialization leads to either cycling between specialist types, or to a population of generalist hosts with a mix of symbiont specialists. If *π_g_* < (*π* + *π*′)/2, the former is the case. If *π_g_* > (*π* + *π*′)/2, the latter is the case.

**Table SI-4.** Summary of stability of equilibria for cooperative specialization. See Table SI-1 for a glossary of these terms.

| Equilibria | Stability |
| --- | --- |
| **Boundary equilibria** |  |
| Matched host and symbiont type: $(0, 0, 1), (1, 1, 0)$. | Always saddle node. |
| Mismatched host and symbiont types: $(0, 1, 0), (1, 0, 1)$. | Always saddle node. |
| Generalist host and monomorphic symbionts: $(0, 0, 0), (1, 0, 0)$. | Always unstable node. |
| **Interior (polymorphic) equilibria** |  |
| Generalist host and polymorphic symbionts: $(u^*, 0, 0)$. | If $\pi_g > (\pi + \pi')/2$ then a stable line.<br>Else, unstable. |
| Coexisting specialist hosts and symbionts: $(1/2, 1/2, 1/2)$. | If $\pi_g > (\pi + \pi')/2$ then unstable. Else, it is a centre. |

##### SI-2.1.1 Boundary equilibria

(0,0,1) and (1,1,0), the matched equilibria, are unstable, since the Λ_1_ > 0. The set of eigenvalues are

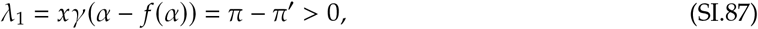

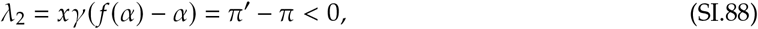

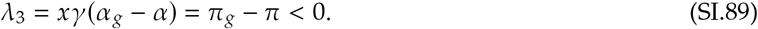

Note that (SI.87) is the opposite of (SI.88) thus, if one case is stable, the other cannot be. A mismatch equilibria exists at (0,1,0) and (1,0,1) at which a host is fixed and its corresponding preferred symbiont is absent. The eigenvalues of the Jacobian matrix evaluated at these equilibria are:

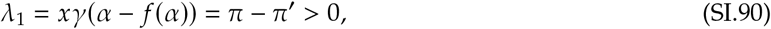

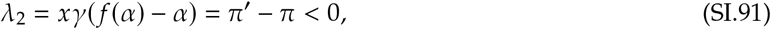

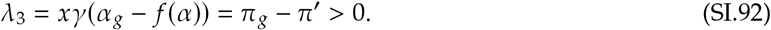

(SI.90) has the opposite sign of (SI.91) and thus this case is also unstable. (0,0,0) and (1,0,0) correspond to a population entirely comprised of generalist hosts and a single type of symbiont. At these equilibria the eigenvalues of the Jacobian are:

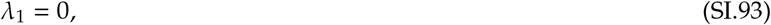

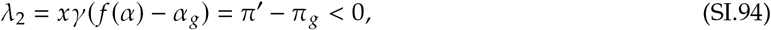

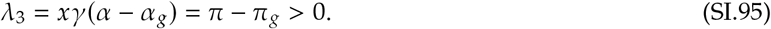

This case is unstable as well. Note that as in the corresponding equilibria under cooperative specialization that if *v*_1_ = *v*_2_ = 0 that *u*\* may once more be any value in [0,1]

##### SI-2.1.2 Internal equilibria

As in cooperative specialization, there is a line of equilibria, *u*\* ∈ [0,1] when 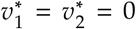. All hosts are generalists, with any combination of symbiont types. We consider the eigenvalues for this line of equilibria:

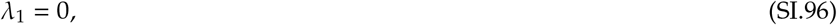

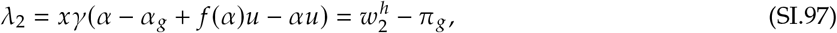

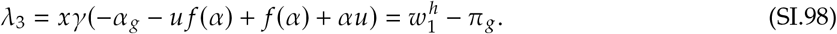

Both (SI.97) and (SI.98) can be negative when (SI.27) is satisfied, i.e. *π_g_* > (*π* + *π*′)/2. Within this interval, the line of equilibria is attracting, while outside of it, it is repelling. We apply the theory of bifurcation without parameters to characterize the dynamics at the endpoints (Liebscher, 2015). Without loss of generality we consider the right end point of (SI.27), by the change of coordinates *μ* = *u* + (*π*_g_ – *π*′)/(*π* - *π*′), *y* = *v*_1_, and *z* = *v*_2_. At the equilibrium (*μ, y, z*) = (0,0,0) the Jacobian is:

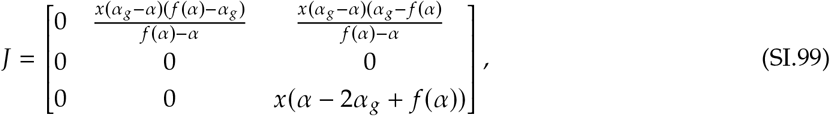

We apply the Center Manifold Theorem to determine the stability of the higher order terms of the system as a function of the bifurcation parameter μ and the centre dimension *y* (*z* = *m*(*μ, y*) = *a*_11_*μy* + *a*_20_*μ*^2^ + *a*_02_ + …). We solve for the coefficients *a_ij_* as before using the invariance condition (SI.29). We find that all coefficients must be zero, thus for this case *m*(*μ, y*) = 0 and dynamics on the centre manifold are thus described by:

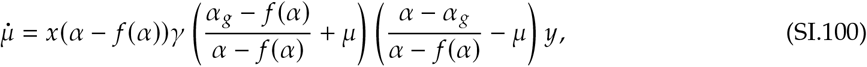

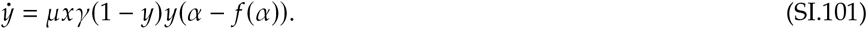

The endpoints of this line of equilibria are stable on the interval *u* ∈ [0,1], thus when *π_g_* > (*π* + *π*′)/2 the generalist equilibrium is stable.

We now consider the internal equilibria for *v_i_* > 0. As before, the host frequency equations (SI.85) and (SI.86) are at equilibrium when (SI.32) is satisfied. From (SI.84) if 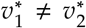 then *u*\* = 0 or *u*\* = 1 i.e. the symbiont population is monomorphic. Without loss of generality, let *u* = 1 giving us

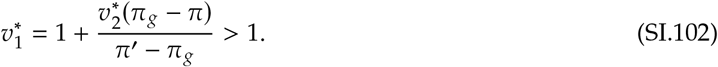

from (SI.32). Thus this type of internal equilibrium is impossible.

We next consider equilibria where 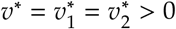. As with cooperative specialization (SI.32), Equation SI.32 with *i* = 1 and *j* = 2 gives us

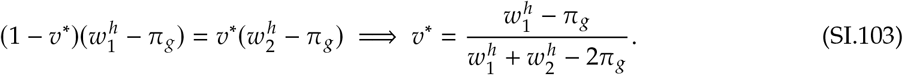

Then, substituting *v*\* into Equation SI.32 with *i* = 2 and *j* = 1, we have

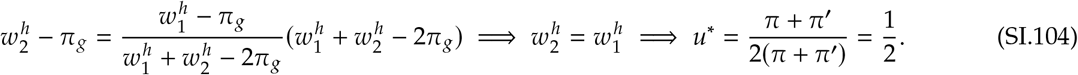

Then, 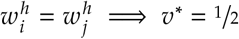 giving us our final polymorphic equilibrium: (1/2, 1/2, 1/2).

The eigenvalues for this case are as follows:

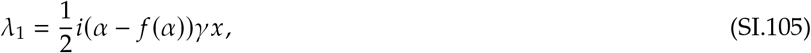

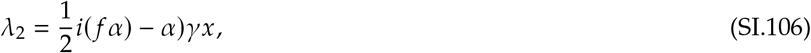

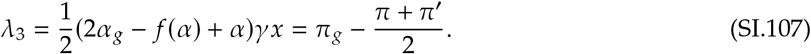

Λ_3_ may be positive or negative, depending on the difference between generalist payoff and specialist average payoff. If it is positive, we have an unstable equilibrium. However, if it is negative, then we can apply the Centre Manifold Theorem to evaluate the nature of the equilibrium.

We change to the coordinate system (*y*_1_, *y*_2_, *z*) with the invertible matrix *P*:

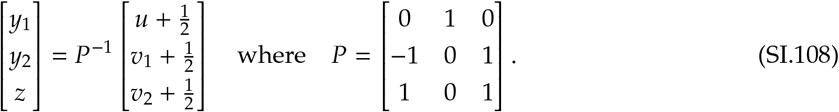

Note that we have shifted the equilibrium to (0, 0, 0). The system in these new coordinates is:

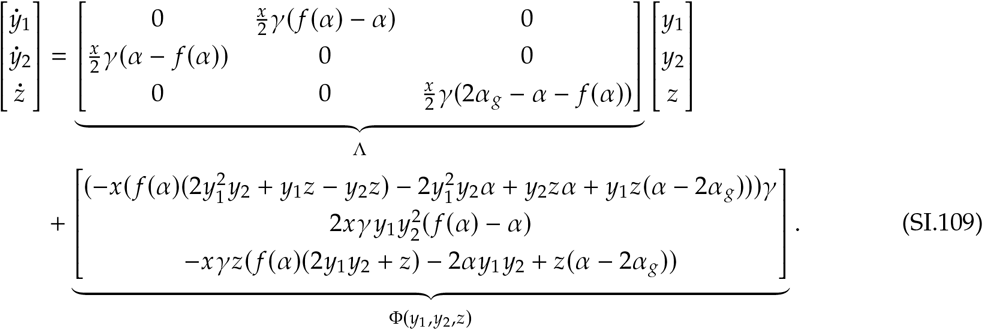

Φ(*y*_1_, *y*_2_, *z*) holds the higher order terms, and Λ = *P*^-1^/*P* is the real Jordan normal form with Jacobian

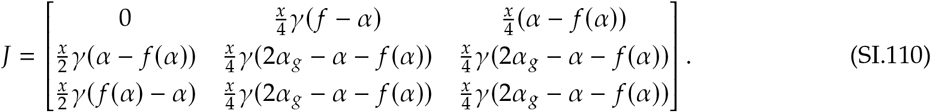

Next we use the Center Manifold Theorem to characterize the stability of the higher order terms of the system as a function of the two center dimensions *y*_1_ and *y*_2_. Setting 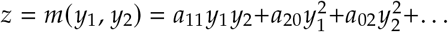 we have the invariance condition

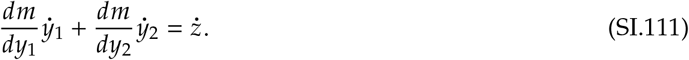

From here, however, we find that all coefficients must be zero and thus *m*(*y*_1_, *y*_2_) = 0. This is because 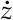 is a multiple of *z*, and thus all terms carry a coefficient of the series. The dynamics on the centre manifold are thus described by

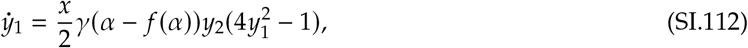

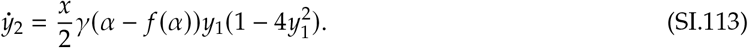

This system has a centre at (0,0). Thus, if *π_g_* < (*π* + *π*′)/2, both specialists can coexist, however this state is a center, and any perturbation away from this node causes specialist hosts to cycle. Numerical simulations of this behavior can be viewed in the supplemental Mathematica notebook `Limit Cycle Simulations.nb` at https://github.com/christopheriancarlson/SpecificMutualism.

#### SI-2.2 Evolutionary invasion analysis

Here we consider evolution of traits *α, γ*, and *x* at the generalist equilibria (SI-2.2.1) and when the population is cycling. In the latter case, we conducted numerical simulations to determine whether (and where along the period of a cycle) a mutant can invade (SI-2.2.2). Table SI-5 summarizes the results.

**Table SI-5.**
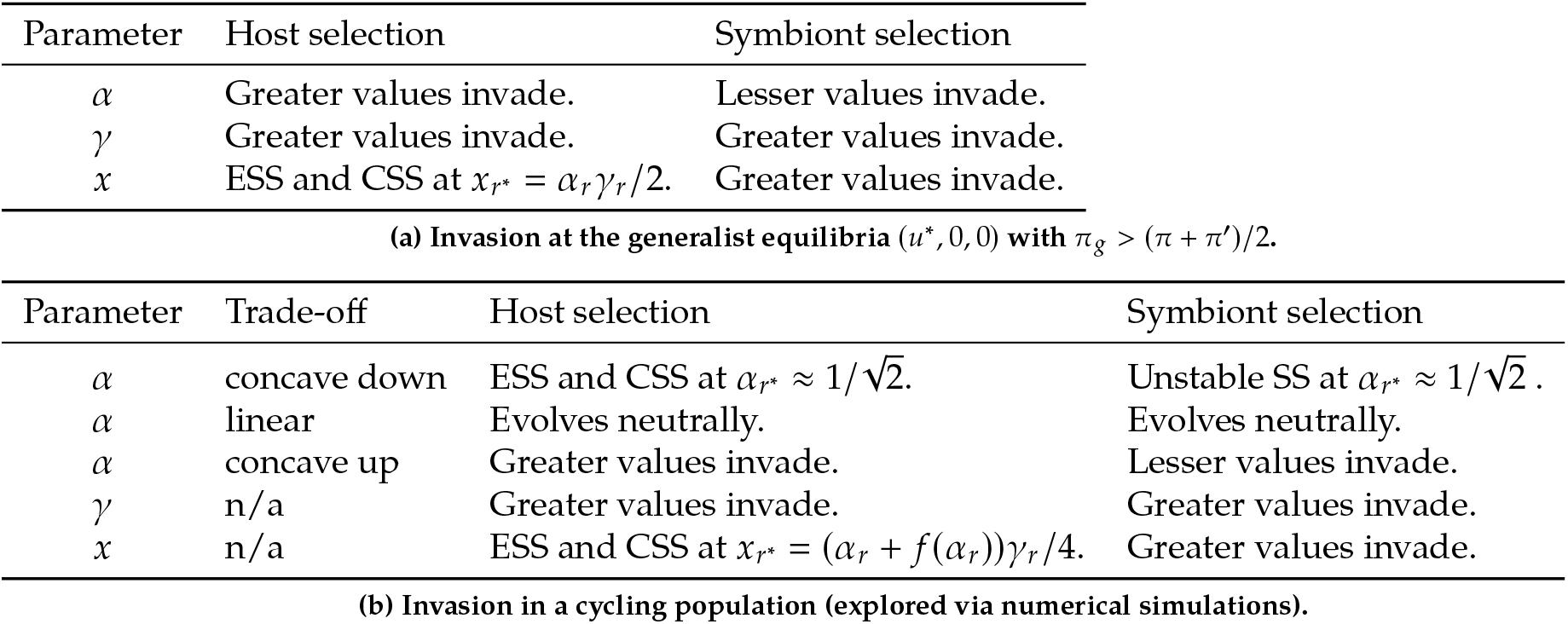
Summary of invasion analyses for antagonistic specialization. SS, ESS, and CSS mean singular strategy, evolutionarily stable singular strategy, and convergence stable singular strategy, respectively.

##### SI-2.2.1 Evolution at generalist equilibria

First, we assume that traits are under selection by hosts at the generalist equilibrium. The initial growth rate of a rare mutant generalist host (whose trait values are once more denoted by the subscript *m*) is:

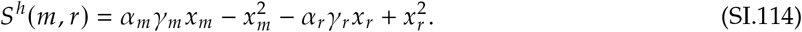

As before, the *m* subscript denotes a mutant trait value, the *r* subscript denotes a resident trait value and *h* denotes that this is the invasion exponent for hosts. Note that unlike SI-2.2 all hosts are generalists, that receive *α_g_* from all symbionts. Therefore, the invasion exponent is not a function of *u*. The selection gradients for *α_m_* and *γ_m_* are:

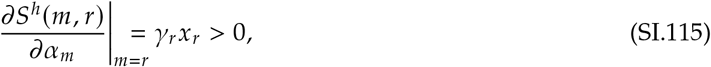

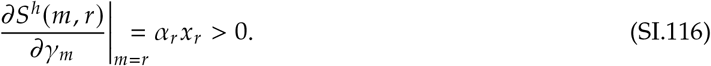

These positive selection gradients indicate that invading host mutants with larger values of either *α* or *γ* will always successfully invade. The *x_m_* selection gradient is

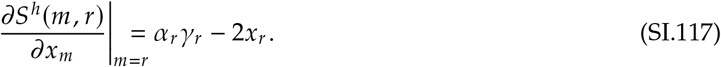

As in cooperative specialization, there is a singular strategy 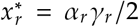. The second derivative at this singular strategy is:

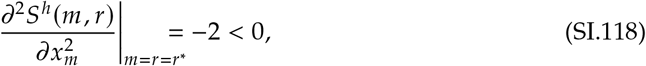

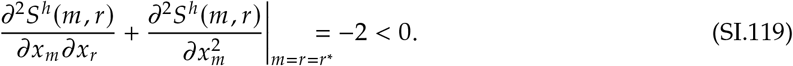

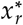 is a local fitness maximum and thus an evolutionarily stable level of host investment. And, it is also convergence stable.

When we assume that the traits are under selection by symbionts at a generalist equilibrium, the invasion exponent for a symbiont that interacts with a generalist host is

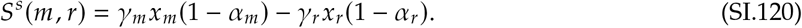

The superscript *s* denotes that this invasion exponent is that of a symbiont. Selection gradients for symbiotic traits are:

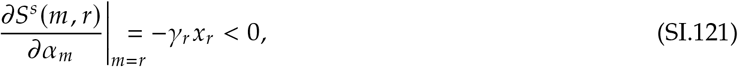

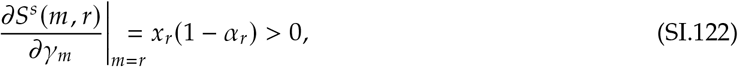

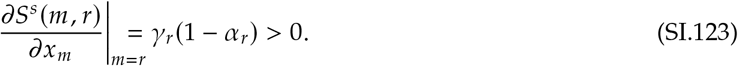

Symbionts with larger values of *γ* and *x* will be favored, while reduced values of *α* will always allow a mutant to invade and fix successfully.

##### SI-2.2.2 Evolution in a cycling population

Here we consider the case in which specialist average payoff exceeds generalist payoff, leading to a cycling population composed of two specialist hosts and two symbionts. Because the frequencies of hosts and symbionts cycle, the timing of mutant invasion influences the success of mutant invasion. Thus, analytical invasion analysis was intractable given that it assumes no periodic variation in strategist frequency. To over-come this analytical limitation we conducted invasion analysis via numerical simulations. In simulations we systematically introduced rare mutants whose trait values differed from those of residents over the course of a resident’s cycle. Because both type 1 and type 2 specialists are present in the cycle, we considered mutants of both types. Generalists are entirely excluded, and are therefore not included in the simulations in SI-4 and 5. We found the frequency of mutants whose trait values differed slightly from those of residents following their introduction and we defined a successful invasion as a case in which the initially rare mutant had increased in frequency long after its introduction. We measured the proportion of the resident’s cycle that could be successfully invaded by the mutant to determine the overall outcome of mutant invasion. These methods are included in the supplemental Mathematica notebook `Antagonistic Invasion Analysis.nb, Antagonistic Parameter simulations X and gamma`, and `Antagonistic Parameter Simulations alpha` which can be found at https://github.com/christopheriancarlson/SpecificMutualism.

**Figure SI-3.**
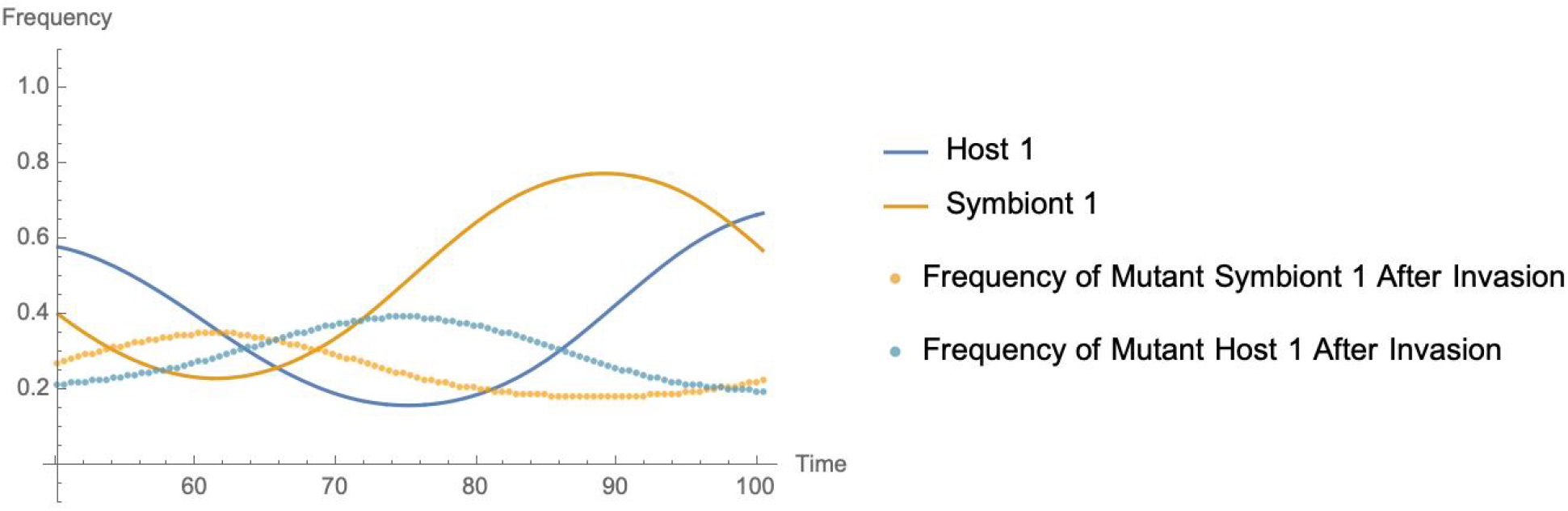
**A sample cycle of hosts and symbionts is shown when** *α* = 0.61, 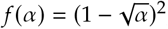, *α_g_* = 0.3, *x* = 0.5 **and** *γ* = 1. **Without loss of generality, we consider type 1 hosts and symbionts. We introduced mutants with a slightly different value of** *α* **from residents systematically along this resident cycle (***α* = 0.605 **for symbionts and** *α* = 0.615 **for hosts). We then measured the average frequency of these mutants long after their introduction from** *t* = 2200 **to** *t* = 2500. **In this figure, mutant frequency is superimposed over the point in the resident cycle at which these mutants were introduced. Mutants invade the most rapidly when the resident of the corresponding type is at the trough of its cycle, e.g. a host 1 mutant replaces a resident host 1 the most quickly when mutant invasion occurs at the trough of the resident host 1 cycle.**

We found that the timing of invasion influenced the degree of success of mutants, which we depict in Figure SI-3. Mutants are most successful when their invasion occurs at a cycle trough of the resident of the corresponding type. They are least successful when the resident of their type is at the peak of its cycle.

Figure SI-4-*c* and Figure SI-4-*d* show that ever increasing values of *γ* will always be favored by selection, and that there is a singular strategy for *x* when *x* ≈ (*α* + *f*(*α*))*γ*/4. Because larger values of *x* will be favored to the left of *x* ≈ (*α* + *f*(*α*))*γ*/4 and smaller values of *x* to the right of *x* ≈ (*α* + *f*(*α*))*γ*/4, we conclude that *x* ≈ (*α* + *f*(*α*))*γ*/4 is both evolutionarily stable and convergence stable. This intermediate value of *x* balances the cost of trade with both a host’s matched and mismatched symbiont. Finally in Figure SI-4-*b* and Figure SI-4 - *d* we find that larger values of *x* wand *γ* will always be favored by symbionts.

**Figure SI-4.**
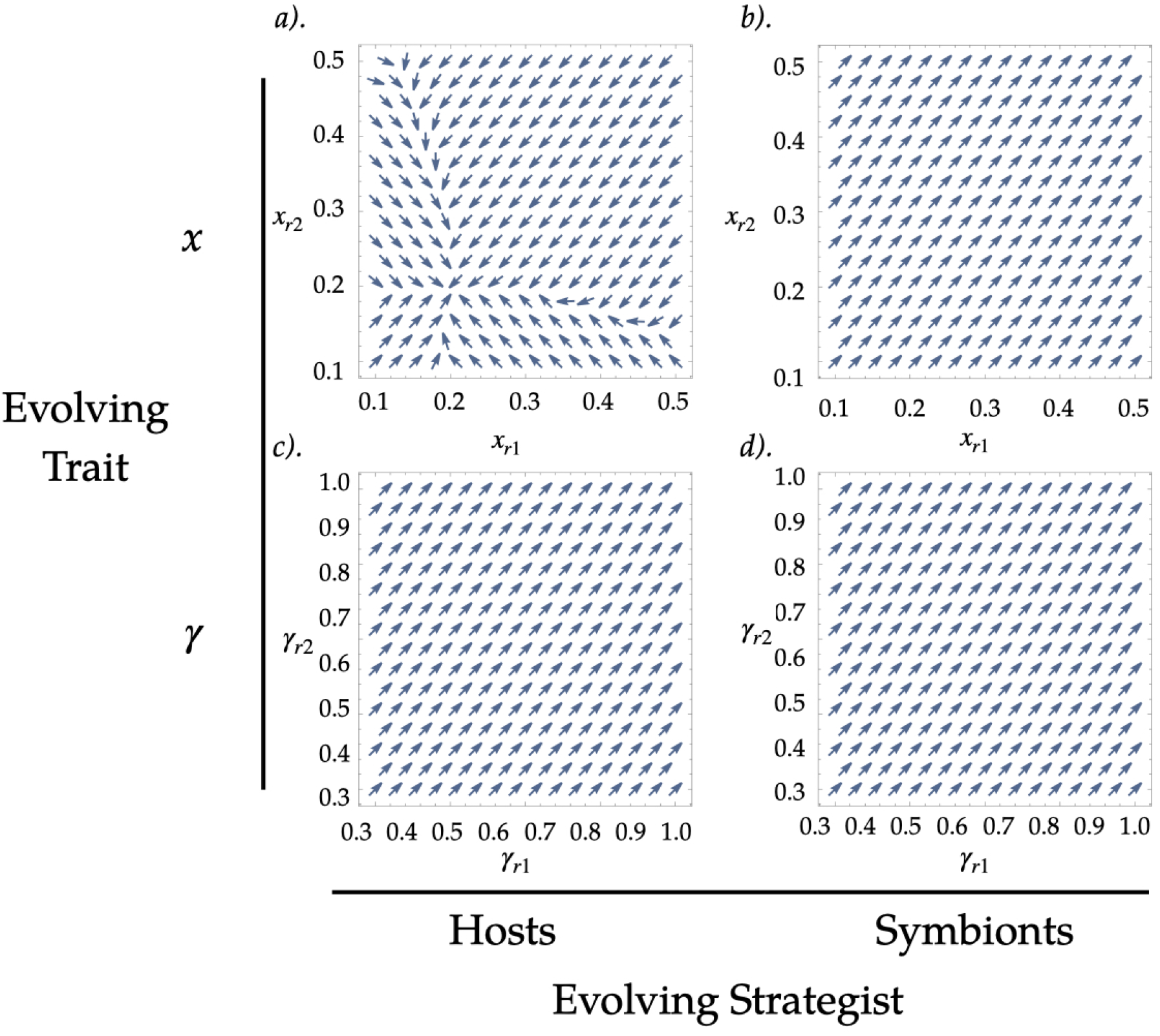
**Vector plots for** *x* **and** *γ* **for hosts and symbionts. In all plots** *α* = 0.8, **and** 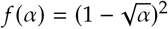. **In plots *a)* and *b*)** *γ* = 1, **in plots *c*) and *d*)** *x* = 0.5. **The** *x* **component of each vector is the proportion of a cycle invaded by a type** 1 **mutant and the** *y* **component is the proportion of the cycle that can be invaded by a type** 2 **mutant. The magnitude of all vectors in all plots is 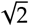.**

## Notes

https://github.com/christopheriancarlson/SpecificMutualism

### Competing Interest Statement

The authors have declared no competing interest.

